# Cancer tolerance to chromosomal instability is driven by Stat1 inactivation *in vivo*

**DOI:** 10.1101/2021.12.03.471107

**Authors:** Michael Schubert, Christy Hong, Laura J. Jilderda, Marta Requesens Rueda, Andréa E. Tijhuis, Judith E. Simon, Petra L. Bakker, Jon L. Cooper, Aristi Damaskou, René Wardenaar, Bjorn Bakker, Sahil Gupta, Anouk van den Brink, Lorena Andrade Ruiz, Miriam H. Koster, Sameh A. Youssef, Danielle Luinenburg, Alex Strong, Thomas Engleitner, Hannes Ponstingl, Gerald de Haan, Alain de Bruin, Roland Rad, Hans W. Nijman, René H. Medema, Marcel A.T.M. van Vugt, Marco de Bruyn, Diana C.J. Spierings, Maria Colomé-Tatché, George S. Vassiliou, Floris Foijer

## Abstract

Chromosomal instability is a hallmark of cancer, but also an instigator of aneuploidy-induced stress, reducing cellular fitness. To better understand how cells with CIN adjust to aneuploidy and adopt a malignant fate *in vivo*, we performed a genome-wide mutagenesis screen in mice. We find that specifically aneuploid tumors inactivate Stat1 signaling in combination with increased Myc activity. By contrast, loss of p53 is common, but not enriched in CIN tumors. Validation in another tissue type confirmed that CIN promotes immune cell infiltration, which is alleviated by Stat1 loss combined with Myc activation, but not with p53 inactivation, or Myc activation alone. Importantly, we find that this mechanism is preserved in human aneuploid cancers. We conclude that aneuploid cancers inactivate Stat1 signaling to circumvent immune surveillance.

## Introduction

Chromosomal instability (CIN) is a phenomenon characterized by frequent chromosome missegregation events, resulting in cells displaying structural and/or chromosomal abnormalities, *i.e.* aneuploidy (*1*). CIN is a hallmark of cancer and associated with tumor recurrence, metastasis and drug resistance and thus poor patient prognosis (*2–5*). However, in untransformed cells, CIN and aneuploidy impair cellular fitness, evidenced by reduced proliferation, decreased transformation potential, a deregulated cellular metabolism, and senescence (*6–8*). This apparent contradiction is also known as the aneuploidy paradox (*9*) and implies that aneuploid cancer cells need to overcome certain barriers to cope with aneuploidy. As aneuploid cancer cells might rely on such coping mechanisms for their survival, targeting these mechanisms could provide a powerful strategy for treating these cancers. However, to develop such therapies, we need to better understand the barriers that aneuploid cells need to overcome to become malignant.

Over the past decade, several aneuploidy-induced stresses have been identified, including proteotoxic stress (*4, 7, 10*), metabolic stress (*8, 11*), and an inflammatory response (*11–15*). Reducing these stresses has been found to improve the overall cellular fitness of aneuploid cells while exacerbating these stresses reduces their fitness. For instance, neuronal cells from individuals with Down syndrome exhibit increased expression of the ubiquitin-mediated proteasome system, and are extremely sensitive to factors that exacerbate proteotoxic stress such as compounds that trigger ER stress (*16*). Similarly, aneuploid yeast cells are extremely sensitive to inhibition protein ubiquitination, as this exacerbates aneuploidy-induced proteotoxic stress (*17*). While such studies have greatly improved our understanding of how individual cells cope with aneuploidy, most were performed in cultured mammalian cells or yeast and therefore cannot fully capture the complete process of tumorigenesis.

To better understand how CIN and aneuploidy lead to cancer, mouse models were generated with mutations that lead to decreased mitotic fidelity (*18, 19*), which revealed that in most genetic backgrounds CIN is a poor instigator of cancer. However, inactivation of p53, a common event in cancer, promotes propagation and transformation of tetraploid cells and the formation of aneuploid tumors (*20*). Indeed, loss of p53 and CIN have been shown to collaborate in tumorigenesis in various models (*12, 21, 22*). However, the exact role of p53 in preventing the outgrowth of aneuploid cells is not fully understood, as for instance expression of a functional p53 does not impair propagation of all aneuploid cells *per se* (*23, 24*).

Complementarily, observational studies of large human cancer cohorts have identified an increased mutational load and decreased leukocyte infiltration in more aneuploid tumors (*25*) Somatic copy number aberrations have been linked to aneuploidy-specific immune evasion, *e.g.* by gaining certain oncogenes or loss of heterozygosity at the HLA locus (*26*). Gene expression signatures have been developed to estimate CIN in human tumors (*27*), but there is some disagreement about the extent they, instead, reflect other processes like CIN-independent proliferation (*28, 29*). Finally, multi-region sequencing efforts have employed tumor heterogeneity to conclusively link oncogene and antigen shuffling to chromosomal instability (*30, 31*). In particular, metastatic samples showed a CIN-driven recurrent amplification of 8q24, encompassing *MYC*, or 11q13, encompassing *CCND1*(*31*).

Both the functional as well as the observational studies have provided valuable insights into how cancer cells cope with stresses introduced by aneuploidy or chromosomal instability. However, to our knowledge there has not been an unbiased and comprehensive comparison of how different cancer drivers enable euploid or aneuploid/CIN transformation into aggressive tumors *in vivo*. Here we present a genome-wide transposon mutagenesis screen in the hematopoietic system of isogenic mice with or without induced CIN. We find that p53 mutations enable both euploid and aneuploid cancer growth, while a combination of Stat1 inactivation and Myc amplification is a driver unique to the CIN phenotype. We further provide strong evidence that this mechanism extends to other cancer types in mice, and is consistent with observations in The Cancer Genome Atlas (TCGA) cohorts.

## Results

An *in vivo* insertional mutagenesis screen using a genetically engineered CIN mouse model Chromosomal instability discriminates cancer cells from healthy cells. Therefore, selectively targeting CIN could provide a powerful strategy to treat many cancers. To develop such therapies, we need to understand which features are unique to CIN cancers. We performed an *in vivo* genetic screen to compare the cancer-driving genes between tumors with and without a CIN phenotype in an otherwise isogenic setting. We took advantage of the fact that CIN alone is a poor instigator, yet a powerful accelerator of cancer when combined with cancer-predisposing mutations (*12, 21, 22, 32, 33*). To limit CIN and mutagenesis to a single tissue, we combined a conditional driver of CIN with a conditional transposon mutagenesis system. For this, we crossed mice harboring our well-characterized conditional Mad2^f/f^ allele (*11, 22*) with mice that carry a cassette of 15 PiggyBac (PB) transposons on chromosome 16 and a Lox-Stop-Lox PB-Transposase, allowing for tissue-specific activation of transposon mutagenesis (*34, 35*). The PB transposable elements in this model can activate or inactivate genes depending on the integration site (**Fig. 1a**) (*34, 35*). We chose to target the hematopoietic system as 1) approximately half of human hematopoietic cancers aneuploid (*36*), allowing us to compare the drivers between CIN and non-CIN tumors and 2) hematopoietic tumors in mice display a relatively short latency. We therefore crossed our mice into an Mx1-Cre background in which Cre-recombinase can be activated in the hematopoietic compartment through PolyI:PolyC (pI:pC) injections (**Fig. 1a**). Two cohorts of mice were treated: 1) 45 Transposon; Mx1-Cre mice and 2) 104 Mad2^f/f^;Transposon;Mx1-Cre mice. Both cohorts were monitored for tumorigenesis alongside two control groups (pI:pC-treated Mad2^f/f^;Mx1-Cre mice and untreated Mad2^f/f^;Transposon;Mx1-Cre mice). Treated Mad2^f/f^;Transposon;Mx1-Cre mice showed a median survival of 13 months compared to 18 months for Transposon;Mx1-Cre mice, confirming that CIN is a powerful accelerator of tumorigenesis, also in this background (**Fig. 1b**). Untreated Mad2^f/f^;Transposon;Mx1-Cre mice developed sporadic hematopoietic malignancies as well, but with longer latencies, consistent with sporadic leaky Mx1-Cre expression. Histology and flow cytometry analysis of the malignancies revealed that about half were poorly differentiated B-cell tumors, and the other half were either of myeloid or T-cell origin (**Fig. S1a-d**, **Fig. 1b**). We quantified Mad2 loss in Mad2^f/f^ tumors by quantitative genomic PCR, RT-PCR and Western blot assays, and confirmed that most Mad2^f/f^ tumors had indeed lost Mad2 expression (**Fig. S1e-i**).

**Fig. 1.**
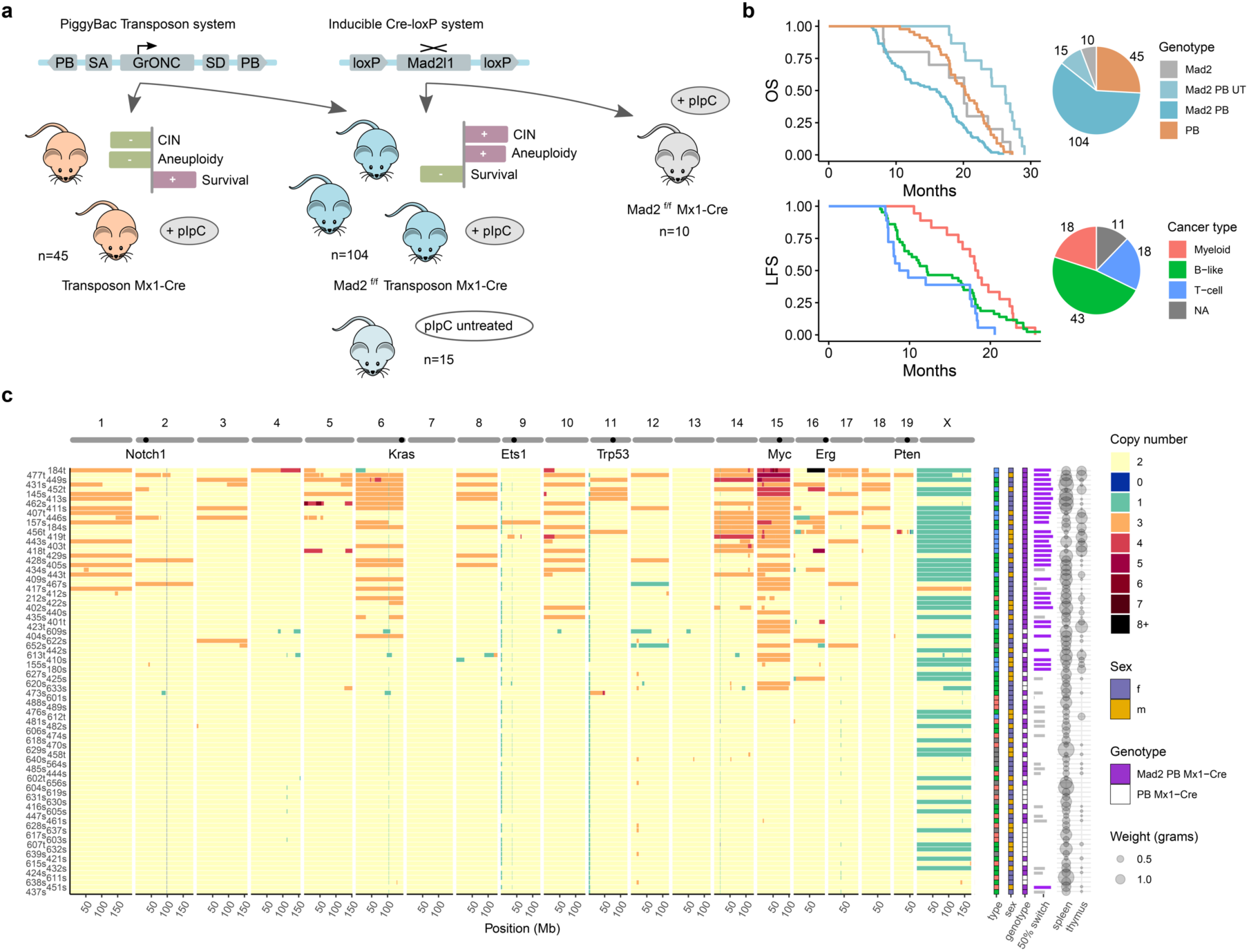
Experimental setup and cohort overview. **(a)** Treatment scheme of mice with either PiggyBac insertional mutagenesis alone or in combination with homozygous Mad2^f/f^ allele. Both genetic modifications are activated via pI:pC. Controls include untreated (UT) mice and Mad2^f/f^ alone. **(b)** Overall survival by genotype (OS, top) or Leukemia free survival (LFS) by consensus tumor type from histology and FACS (bottom). Combined Mad2^f/f^ PB constitutes the largest part of the cohort, followed by PB-only. The combined genotype is the most aggressive and untreated mice show the longest survival. Both individual genotypes fall in between. For tumor types, T-ALLs are the most aggressive, followed by B-like tumors and myeloid tumor bearing mice survive the longest. Tumor types are enriched in the respective genotypes. **(c)** Average DNA copy number patterns across the whole cohort, including tumor type, sex, genotype, switching efficiency, and tumor weight in both spleen and thymus. Chromosome 15 is the most amplified, followed by chromosome 6. T-ALLs are the most aneuploid, myeloid the least. B-like tumors span the whole range. Aneuploidy is strongly correlated with the Mad2^f/f^ genotype, most tumors of which are also homozygously switched. Unswitched tumors are markedly more euploid, with a single exception of a switched tumor. Heterozygous switching does not confer CIN (grey bars).

To quantify aneuploidy, we performed shallow whole genome sequencing for all tumors, which showed that all Mad2^f/f^ tumors, except one, and only a few Transposon-only tumors were aneuploid (**Fig. 1c**). As expected, tumors with only heterozygous inactivation of Mad2 were not affected by the CIN phenotype. In terms of whole-chromosome copy number changes, we find that chromosome 15 (harboring *Myc*) was the most frequently amplified in the CIN-high tumors, and the only chromosome with recurrent amplifications of more than one additional copy. The next most frequently amplified was chromosome 6, harboring *Kras*. Both *Myc* and *Kras* are potent oncogenic drivers, supporting the hypothesis that CIN enables tumors to shuffle their genome content to select clones with a favorable oncogene to tumor-suppressor gene ratio (*26, 37*). We also observe recurrent amplifications of the region of chromosome 16 that harbors *Erg*, a well-known driver of leukemias (*38*). Stratification per tumor type revealed that myeloid malignancies were rarely aneuploid, and that T-ALLs on average were more aneuploid than B-cell precursor malignancies.

### Transposon insertions and RNA-seq reveal a CIN-dependent Stat1/Myc axis

We sequenced the transposon integration sites and mapped Common Insertion Sites (CIS), *i.e.* transposon integrations that affected a gene or region more often than would be expected by chance, across the entire mouse cohort (**Fig. 2a**). Among the most affected genes were *Trp53*, *Crebbp*, and the known hematopoietic drivers *Ets1*, *Erg*, *Ikzf1*, *Foxn3*, *Pten*, and *Runx1* (**Fig. S2a**), consistent with a previous hematopoietic screen (*35*). We also quantified whether a CIS is CIN-specific by testing for a difference in aneuploidy in inserted *vs.* non-inserted samples as aneuploidy correlates well with karyotype heterogeneity in single-cell DNA sequencing, an established measure for CIN (**Fig. 2b**). Among the most significantly enriched genes were *Erg*, *Hs6st3*, *Dnm3*, and *Stat1* (**Fig. S2b**). We further used a subnetwork identification algorithm on protein interactions to narrow down our list of hits to the most important altered biological processes. We retained most of our top hits, suggesting they may target a common process (**Fig. S2c-d**). Ranking the cleaned-up list by degree of aneuploidy, we found *Erg* and *Stat1* as the most CIN-specific hits, *Ets1* and *Trp53* as non-specific, and *Pias1* as the most euploid-specific hit (**Fig. 2c**). Interestingly, Pias1 is a known inhibitor of Stat1(*39*), also by name (Protein inhibitor of activated Stat1). This, together with the fact that Stat1 is central to both the cohort and the CIN-specific subnetwork (**Fig. 2d**, **Fig. S2c-d**), leads us to conclude that inactivation of Stat1 is likely the main requirement for hematopoietic tumors to tolerate CIN. Interestingly, insertions involving Stat1 interactors were generally enriched in euploid tumors, the opposite of Stat1 itself (**Fig. S2d**), suggesting that they may also act as inhibitors.

**Fig. 2.**
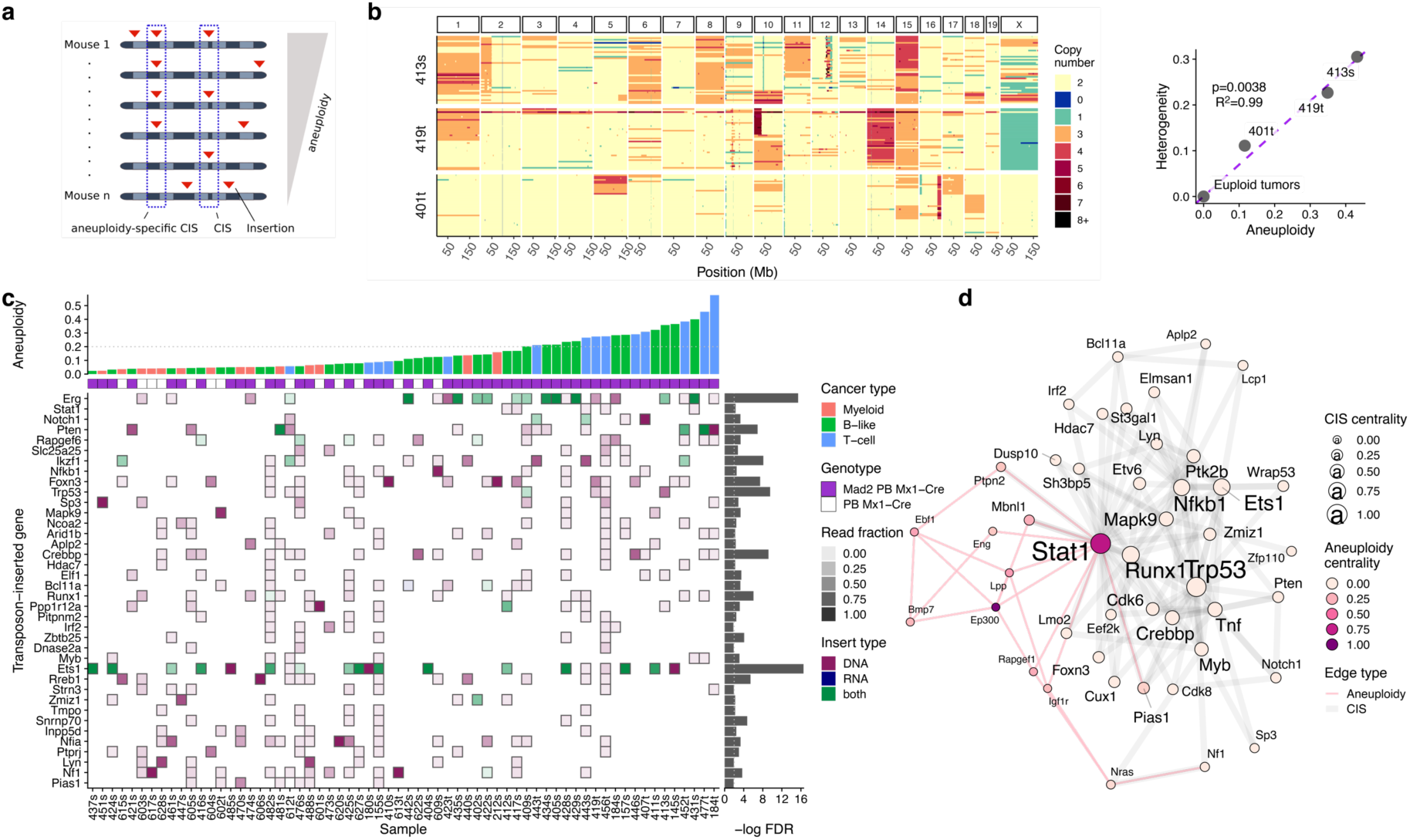
Common Insertion Sites (CIS) in the mouse transposon cohort. **(a)** Schema of finding CIS that are occurring more than expected by chance, or in addition occurring more towards either end of the aneuploidy spectrum. **(b)** Single-cell whole genome sequencing of three tumors shows an almost-perfect correlation between aneuploidy and copy number heterogeneity, a measure of CIN. It is the basis of treating aneuploidy and chromosomal instability interchangeably in the analysis of our mouse cohort. **(c)** Overview of the CIS. Mice are ordered from the most euploid to most aneuploid from left to right, insertion sites from the most euploid to most aneuploid from bottom to top. The color indicates evidence for the insertion found from DNA or RNA, with the shade corresponding to the relative DNA read count number. *Ets1* and *Erg* are the strongest general CIS, with *Erg* and *Stat1* the most aneuploidy-enriched, and *Pias1* the most euploid-enriched. **(d)** Core of the merged insertion subnetwork of both general (grey lines) and aneuploidy-specific CIS (pink lines). *Trp53* is central in the general network, *Ep300* in the CIN network, and *Stat1* in both. All tests performed using a linear model unless stated otherwise.

To better understand the biology of the euploid and aneuploid malignancies, we next analyzed the transcriptomes of all tumors. We confirmed our earlier FACS-based cancer type classification with the gene expression of differentiation markers. Samples clustered according to their cancer type (**Fig. 3a**), and showed an upregulation of the respective lineage (**Fig. 3b, S3a**), *e.g.* CD3 for T-ALLs, Ebf1 for B-like precursors, and Ly6g (GR-1)/Lyz1 for myeloid malignancies. In addition, B-like precursors showed a higher expression of cKit, and a subset harbored an aberrant activation of the Ets1 or Erg transcription factors (B220 low in FACS; **Fig. S1c**), similar to a human leukemia cohort (**Fig. S3b**). Interestingly, the level of aneuploidy in our mouse cohort also reflects that in human pro/pre-B-ALL (Ebf1 and Ets1), and hyperdiploid ALL (Erg). T-ALLs and myeloid leukemias can be both euploid and aneuploid, although there is a bias for more aneuploid T-ALLs and more euploid myeloid leukemias in our mice compared to the human cohort (**Fig. S3c**).

**Fig. 3.**
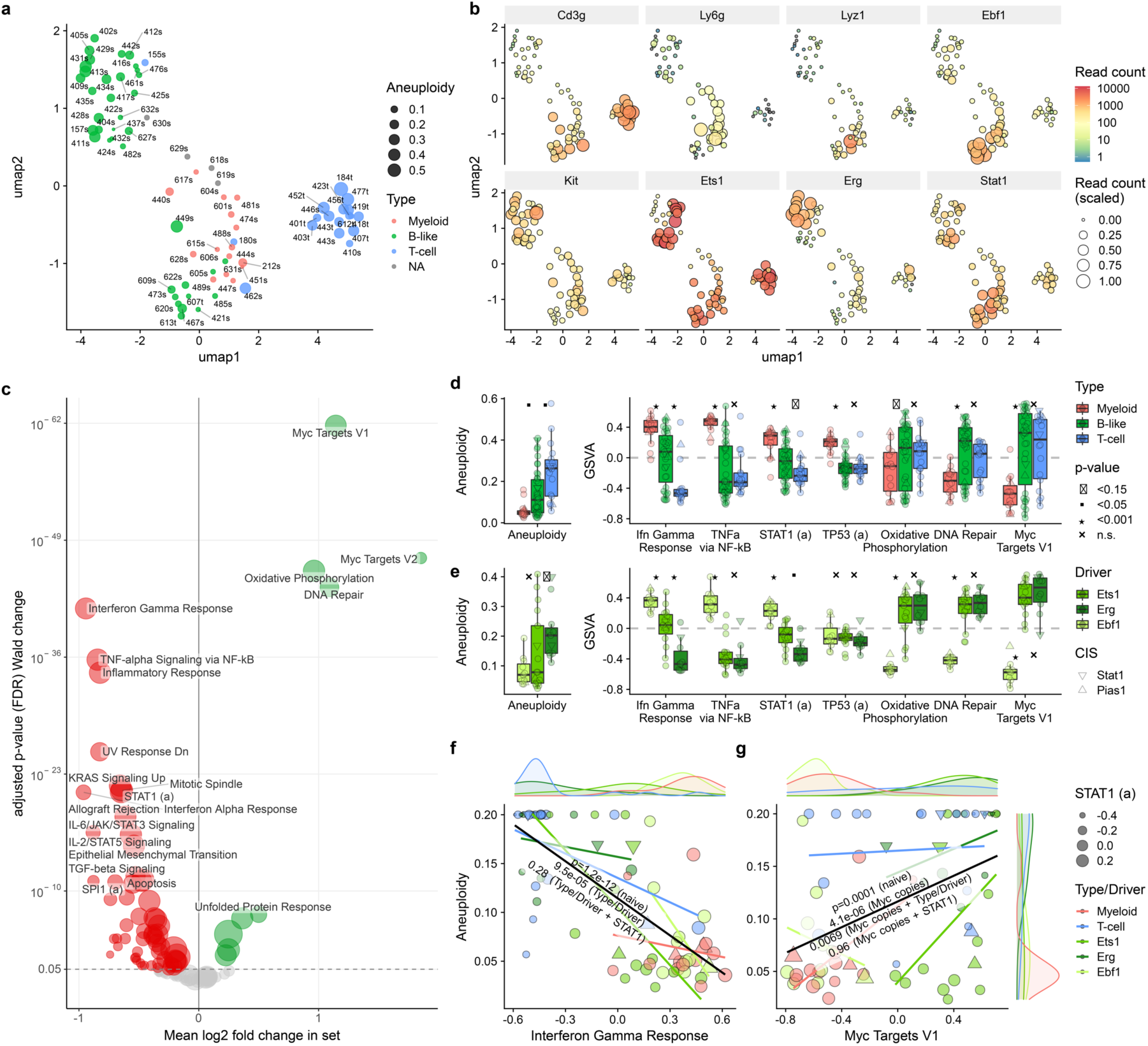
Gene expression in the mouse transposon cohort. **(a)** Dimensionality reduction (UMAP) plot of the gene expression of known differentiation markers shows a distinct cluster for T-ALLs and Ets1/Erg-driven B-like tumors. Ebf1-driven B-like and myeloid tumors show a continuum, in agreement with the histiocytic sarcoma transdifferentiation identified by histology and FACS markers. **(b)** Gene expression of markers for the tumors shows expected expression of Cd3 and T-ALL, Ets1/Erg/Ebf1 in B-like, and Ly6g/Lyz in myeloid tumors. Kit expression is consistent with Ets1/Erg locking tumors in a precursor state that has low Stat1 expression, and Stat1 expression is consistent with baseline expression of the interferon response. **(c)** Differential expression with aneuploidy, corrected for tumor type, shows a downregulation of the Interferon Response and Stat1 target genes, and upregulation of the Myc Targets Hallmark gene set. Oxidative Phosphorylation and DNA repair sets are also upregulated. **(d)** Aneuploidy levels and gene set expression between tumors mirrors the effect observed within tumors, also **(e)** when splitting B-like tumors into its specific drivers. **(f)** Sample-level expression of the Interferon Gamma Response is strongly correlated with aneuploidy, which can be fully explained by tumor subtype in combination with Stat1 activity, but not subtype alone. **(g)** Myc targets positively correlate with aneuploidy, which can be explained by Myc DNA copies only in combination with Stat1 activity, but not by Myc copy number or subtype alone (parenthesis indicate covariate in regression). Boxplots show median and quartiles, whiskers the interquartile range. All tests performed using a linear model unless stated otherwise. Transcription factor activity inference indicated with uppercase letters and (a) for activity.

Next, we quantified which gene expression changes correlate with increased levels of aneuploidy or CIN in our cohort. While correcting for potential confounders of different cancer types, we find an overall strong signal for an inactivation of Stat1 and the interferon/inflammatory response, and an upregulation of Myc targets, with Oxidative Phosphorylation and DNA repair as second-line hits (**Fig. 3c**). The p53 response and the expression of p53 transcriptional targets were largely unchanged, in agreement with our observation that *Trp53* did not show preferential transposon insertions in either euploid or aneuploid tumors. If we, instead of correcting for different cancer types, use their respective gene set expression, we find that the observed differences mirror the different aneuploidy levels also across cancer types (**Fig. 3d**). Splitting our B-like cancer type into Ebf1, Ets1, and Erg-driven tumors, we again observe a consistent pattern between differential expression and aneuploidy/CIN levels (**Fig. 3e**).

While Stat1 is thought of as the primary driver for the interferon response (*40*), we have yet to show that this is true for our cohort. To do this, we inferred Stat1 activity from expression of its target genes, and tested whether this activity can fully explain the correlation between CIN and the expression of the interferon response gene set (**Fig. 3f**). Indeed, we find that when conditioning the association on the cancer type and Stat1 activity, this correlation drops from strongly significant to not significant, indicating that the interferon response is likely driven via Stat1. Similarly, we can show that the correlation between Myc targets and CIN can be explained by Myc copies in combination with Stat1 activity, but not by Myc copies alone or in combination with cancer type (**Fig. 3g**), indicating that Stat1 restricts Myc activity. We conclude that, in our model system, aneuploid hematopoietic malignancies inactivate inflammatory signaling through Stat1 inactivation. CIN, on the other hand, enables oncogene amplification, which in our model predominantly involved Myc. Stat1 inactivation in combination with higher Myc levels yield a high transcription level of Myc target genes.

### Transformed 3T3 cells demonstrate Stat1-driven immune recruitment and immune cell activation in response to CIN

We next set out to validate whether our identified driver genes indeed promote malignant transformation of CIN cells *in vivo*. For this, we engineered an untransformed mouse fibroblast cell line (NIH/3T3) with the main factors that we identify to selectively drive tumors with a CIN phenotype (Myc overexpression with or without Stat1 inactivation). As a non-CIN specific control for transformation, we also engineered p53-deficient 3T3 cells with or without Stat1 (**Fig. 4a**, **Fig. S4a-c**). We chose 3T3 fibroblasts as they represent a very different tissue type and thus allow us to generalize our findings. Furthermore, 3T3 cells can be allografted in immuno-proficient mice to study the interaction of the engineered cancer cells with a fully functional immune environment. To induce chromosomal instability, we introduced dnMCAK, a mutant version of the tubulin-binding protein MCAK (also known as KIF2C) into all four 3T3 genotypes (Myc^OEX^ or p53^KO^, each with or without Stat1). As a non-CIN control, we in parallel introduced wild type KIF2C (*41*) (**Fig. 4a**). CIN phenotypes were confirmed by time-lapse microscopy imaging (**Fig. S4d-f**) and caused a moderate proliferation increase for Myc^OEX^;Stat1^KO^ and decrease for p53^KO^, albeit not significantly (**Fig. 4b**).

**Fig. 4.**
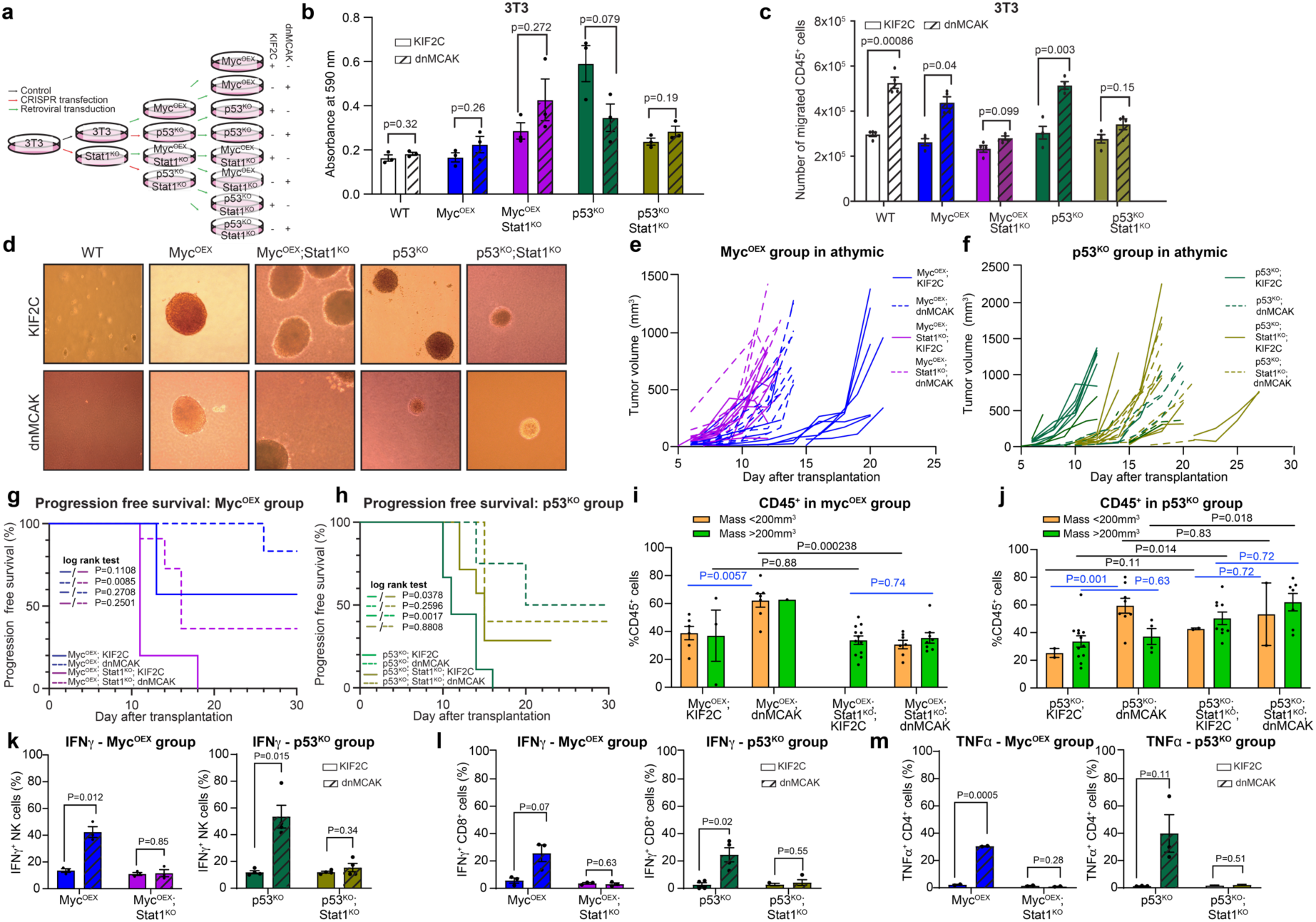
Chromosomal instability promotes Stat1-mediated inflammatory signaling and immune recruitment. **(a)** Schematic overview depicting steps to generate genetically engineered 3T3 transformed lines. **(b)** Crystal violet assay showing the difference in the growth of the engineered 3T3 cell lines with KIF2C (CIN^low^) and dnMCAK (CIN^high^). **(c)** A transwell migration assay demonstrated increased immune cell attraction towards 3T3 cells with a CIN phenotype in a Stat1 proficient background. Inactivation of Stat1 attenuated this effect. **(d)** Representative images from 3T3 genotypes grown in soft agar as an *in vitro* readout of malignant transformation of the engineered 3T3 cell lines. **(e,f)** Inoculation of immunocompromised mice with the 3T3 engineered cell lines overexpressing Myc^OEX^ **(e)** or with a p53^KO^ background **(f)** shows similar growth kinetics *in vivo* compared to *in vitro* (see panel **b**) without a significant growth penalty imposed by CIN. **(g-h)** Progression free survival (tumor size < 500 mm^3^) for tumor fragments allografted in both flanks of immunocompetent Balb/c mice from Myc^OEX^ **(g)** and p53^KO^ **(h)** 3T3 cell lines with or without Stat1 inactivation (n=8 mice per genotype). CIN increased tumor latency for Stat1 proficient tumors and Stat1 loss reduced tumor latency. **(i-j)** Quantification of the CD45^+^ immune cell fraction by flow cytometry within the allograft tumors showed an increase in Myc^OEX^ **(i)** and p53^KO^ **(j)** tumors, which was rescued by Stat1 inactivation in Myc^OEX^ tumors, but not in p53^KO^ tumors. **(k-m)** Cytokine production measured by FACS as a readout of *ex vivo* immune cell activation in NK cells **(k)**, CD8^+^ cells **(l)** or CD4^+^ cells **(m)** co-cultured with Myc^OEX^ (left panels) and p53^KO^ 3T3 cells (right panels) reveals that 3T3 cells that display CIN activate each of these immune cell types and that Stat1 inactivation alleviates this. Error bars represent mean ± standard error of mean (s.e.m). **(b,d,i-m)** Significance tested using a two-sided t-test and **(g-h)** log rank test.

Given the established roles of Stat1 and Myc in inflammation and immune attraction (*42–44*), we then quantified migration of CD45^+^ immune cells (primary isolated Balb/c splenocytes) towards our engineered 3T3 genotypes in a trans-well assay. We find that dnMCAK-induced CIN promotes immune cell migration, and that this is mildly reduced upon overexpression of Myc, but not upon p53 loss. This migration is largely prevented by concomitant Stat1 inactivation in both Myc^OEX^ and p53^KO^ cells (**Fig. 4c**), suggesting that Stat1 activity drives CIN-dependent immune cell infiltration. We next assessed anchorage independent growth as a readout for transformation, which confirmed that all eight engineered genotypes (**Fig. 4a**) grow in soft agar, unlike the parental cell line (**Fig. 4d**). While Myc^OEX^ cells formed large colonies irrespective of other genetic modifications, p53^KO^ cells displayed inhibited growth with either CIN and Stat1^KO^, or the combination thereof. We then inoculated immune-compromised mice with the engineered 3T3 cell lines and found that all cell lines formed tumors efficiently within 13-20 days. Myc^OEX^;KIF2C tumors showed a higher latency compared to either dnMCAK or Stat1^KO^, with the combination being the most proliferative (**Fig. 4e**). By contrast, p53^KO^;KIF2C were the fastest to grow out, with a delay upon either CIN or Stat1^KO^ (**Fig. 4f**), in agreement with their *in vitro* growth characteristics.

To determine the impact of a functional immune system on tumor development, we next allografted 3T3 tumor fragments into Balb/c mice and followed them during tumor progression. Unlike in immuno-compromised mice, we find that Myc^OEX^;dnMCAK cells do not show progressive tumor growth (**Fig. 4g, Fig. S5a**). This suggests that they proliferate efficiently themselves but are eliminated by a functional immune system. Concomitant Stat1^KO^ rescues this CIN-imposed growth delay and restores progressive tumor growth in these mice. By contrast, p53^KO^ tumors generally progress but show a significant delay with dnMCAK compared to non-CIN cells (**Fig. 4h, Fig. S5b**). This difference is compensated by concomitant Stat1^KO^, but these genotypes are still not as aggressive as p53^KO^;KIF2C.

We next quantified immune cell infiltration for all transplanted tumor fragments, including tumors that showed little to no growth (<200 mm^3^). We find that dnMCAK-induced CIN promotes immune infiltration (*i.e.* the fraction of CD45^+^ cells) in Myc^OEX^ tumors by ∼50%, which is rescued by Stat1 loss (**Fig. 4i, Fig. S5d-e**). By contrast, in p53^KO^ tumors, dnMCAK and Stat1^KO^ tend to increase the fraction of CD45^+^ cells, despite efficient outgrowth of these tumors (**Fig. 4j, Fig. S5d-e)**. These observations suggest that CIN impairs Myc^OEX^ tumor outgrowth by recruiting immune cells that Stat1 loss can rescue, while p53^KO^ tumors are less influenced by immune infiltration. Further characterization of the immune cell types in Stat1 proficient tumors revealed that dnMCAK-induced CIN increases the fraction of CD4^+^, CD8^+^, NK and CD19^+^ cells within the CD45^+^ population, particularly in tumors that showed little growth (<200 mm^3^, **Fig. S5f-i**). Inactivation of Stat1 does not change the immune landscape drastically, except for decreasing the fraction of NK and CD19^+^ cells in CIN-driven Myc^OEX^ tumors and decreasing the fraction of CD19^+^ in p53^KO^ tumors (**Fig. S5f-i**). To determine the functional importance of the Stat1 loss-imposed alteration in the *in vivo* immune landscape, we isolated splenocytes from Balb/c mice and exposed them to all eight engineered 3T3 cell lines. We quantified immune cell activation by measuring IFNγ production for NK and CD8^+^ cells and TNFα production for CD4^+^ cells. We find, in agreement with work of others (*13, 15, 45, 46*), that CIN activates NK, CD4^+^, and CD8^+^ cells. Furthermore, we find that Stat1 inactivation in the 3T3 cells alleviates this response, both in a Myc^OEX^ and p53^KO^ background (**Fig. 4k-m**). From these *ex vivo* experiments, we conclude that Stat1 is required to activate NK, CD4^+^ and CD8^+^ cells in tumors that display CIN.

Altogether, these data show that loss of Stat1 activity is required for the *in vivo* growth of Myc-driven tumors with a CIN phenotype. By contrast, p53-deficient tumors grow most efficiently in a non-CIN setting, irrespective of the presence of an immune system. Importantly, these findings functionally validate the significance of Stat1 inactivation from our transposon screen.

### STAT1/MYC drive CIN tolerance in human cancer

Having shown that Myc activation necessitates Stat1 inactivation in chromosomally unstable mouse tumors, we were wondering if the same holds true in human cancers. For this, we took 11 TCGA cohorts for which a separate quantification of immune and stromal contamination was available (ESTIMATE algorithm, details methods). We quantified the expression change of both MYC target genes and STAT1 transcription factor target genes with aneuploidy, and found that most cancers activate MYC targets and inactivate STAT1 target genes (**Fig. 5a**). We further confirmed this by showing that aneuploid tumors show a higher fraction of STAT1 mutations than euploid tumors (**Fig. 5b**), in agreement with the transposon insertion data from our hematopoietic mouse tumors. However, STAT1 mutations only rarely occur in human cancers (2.3%), so there must be additional aberrations to explain the widespread inactivation we observed in our transcription factor activity inference (**Fig. 5a**). One such previously identified factor is a recurrent loss of the interferon alpha gene cluster on chromosome 9, often in combination with *CDKN2A* (*47*). Still, the combined prevalence of both alterations only amounts to 8% across human tumors. This difference may be explained by STAT1 interactor mutations in both our mouse cohort (**Fig. S2d**) and human cancers (**Fig. 5c**), but we need additional data to estimate the effect that each of these mutations may have.

**Fig. 5.**
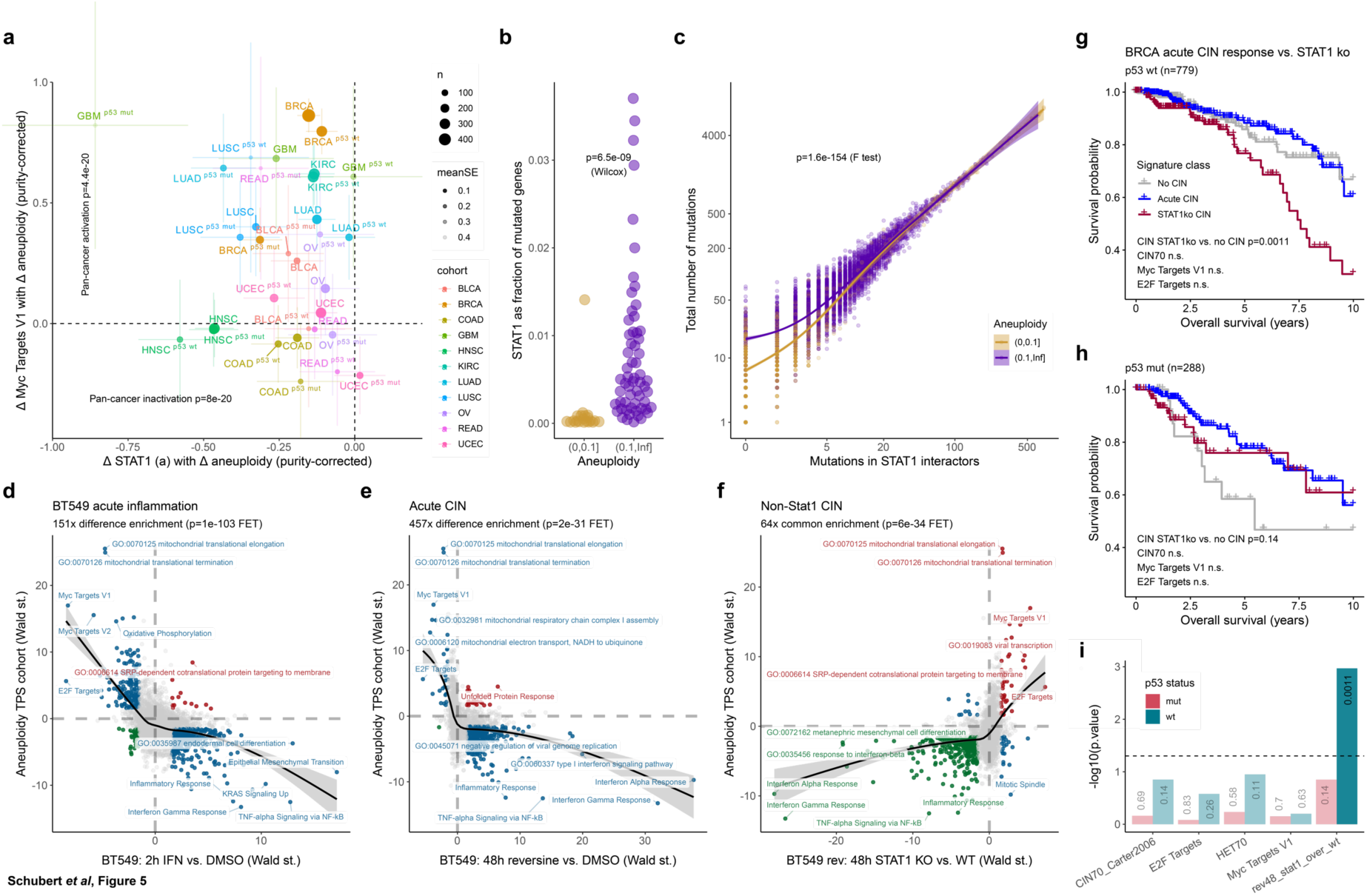
STAT1 in human aneuploid cancer. **(a)** Across 11 TCGA cohorts, all show decrease of STAT1 target genes with increasing levels of aneuploidy when regressing out tumor purity. Myc targets are increased for most cohorts. *TP53* mutated and wild-type samples show the same trend. Bars represent standard error of the slope. **(b)** *STAT1* mutations make up a higher fraction of total mutations in aneuploid samples compared to euploid samples. **(c)** STAT1 interactor mutations make up a lower fraction of total mutations in aneuploid compared to euploid samples when the total mutation rate is low, consistent with transposon insertions. **(d)** Interferon and reversine **(e)** treatment of BT549 cells show the opposite gene expression difference to the CIN adaptation in our mouse transposon cohort for most gene sets in a merged collection of MSigDB Hallmarks, DoRothEA and Gene Ontology. **(f)** Reversine-treated BT549 cells with *STAT1* knockout compared to wild-type show very similar expression changes to CIN adaptation in the transposon cohort. FET Fisher’s Exact Test of the four colored quadrants. **(g)** Cox Proportional Hazards model shows a difference in TCGA breast cancer 10-year overall patient survival when the STAT1^ko^ signature is present compared to when it is absent for p53 wild-type tumors, but not for p53-mutant tumors **(h)**. Alternative explanations like proliferation and CIN signatures show no significant difference **(i)**. Cox models regressed out age at diagnosis and tumor purity. All tests performed using a linear model unless stated otherwise.

In order to address this, we utilize gene expression changes upon STAT1 inactivation in a CIN setting. We chose the chromosomally stable BT549 breast cancer cell line to model a tissue with high MYC activation and on average only moderate STAT1 downregulation. We treated BT549 cells with either Type I interferon or reversine, an MPS1 inhibitor that induces CIN (**Fig. S6a**). RNA sequencing confirmed a transcriptional inflammatory response common to both treatments, and revealed an overall opposite effect compared to the acquired CIN tolerance in our mouse cohort (**Fig. 5d-e**). We then removed *STAT1* via CRISPR-mediated gene knockout and found that the gene expression changes in a CIN setting strongly mimicked the acquired CIN tolerance of our mouse cohort (**Fig. 5f**). Phenotypically, the knockout did not significantly alter proliferation or CIN (**Fig. S6b-d**), but did promote survival and decreased immune cell attraction and activation in response to CIN (**Fig. S6e-h**), consistent with another cell line (**Fig. S6i-m**) and our transformed mouse fibroblasts (**Fig. 4**). This suggests that STAT1 inactivation is a conserved mechanism for CIN tolerance across cell types and species.

Importantly, the latter experiment provided us with a tool to score STAT1 inactivation in a CIN setting across the TCGA breast cancer cohort independent of known markers such as a *STAT1* mutation or *IFNA* loss. We confirmed that our scores positively correlate with aneuploidy and previously published CIN signatures (**Fig. S7a**) as orthogonal measures of CIN. If they truly reflect CIN levels, we would expect them to also correlate with the clinical outcome of CIN, such as treatment response or metastasis formation. We hence divided the human breast cancer patients into CIN-low (no inflammation, no STAT1^KO^ expression signature), acute CIN (high inflammatory signature) and STAT1^KO^ CIN (no inflammation, STAT1^KO^ signature). Indeed, we observed a significant decrease in 10-year overall survival in the STAT1^KO^ patients compared to inflammatory CIN or no-CIN for *TP53* wild-type tumors (**Fig. 5g**). This difference was not present in *TP53* mutant tumors (**Fig. 5h**), and could neither be explained by proliferation (E2F targets), Myc target genes, or other available CIN signatures (**Fig. 5i**).

While it seems counterintuitive that p53-mutant tumors show a better patient survival effect when the STAT1^KO^ signature is present (**Fig. 5h**) contrary to p53 wild-type tumors (**Fig. 5g**), this observation is consistent with our 3T3 data. Murine p53-driven tumors (**Fig. 4j**) showed a higher immune infiltration than Myc-driven tumors (**Fig. 4i**), even when Stat1 was inactivated, and a delayed outgrowth in their primary host adaptation (**Fig. 4f**). As human STAT1^KO^ breast cancer p53 mutant tumors also show a higher immune infiltration than their p53 wild type counterparts (**Fig. S7b**), the better survival of patients with p53 mutant cancers without STAT1 signaling can be explained by an increased immune surveillance. This leads us to the conclusion that MYC activation co-operates with STAT1 inactivation for immune evasion across cancer types and species boundaries, where STAT1 inactivation allows for increased MYC target gene expression (**Fig. S7c**) that cannot be explained by alternate factors (**Fig. S7d**).

## Discussion

We have presented an *in vivo* CIN screen that, from a common starting point of a genetically engineered mouse model, mirrored random gene activation or inactivation using genome-wide transposon insertional mutagenesis. The advantage of this approach is that it takes into account co-evolutionary trajectories between oncogene-driven carcinogenesis and the response of the immune system that is not modeled in the classic yeast or cell culture studies. Furthermore, the isogenic starting point offers controlled conditions with a limited number of potential external factors that may influence this process, unlike observational studies of a human cancer cohort. Owing to this, we can make a well-founded statement about the relative importance of different kinds of oncogenes in an engineered CIN *vs.* non-CIN setting.

A particularly interesting aspect of our findings is that, while p53 has often been described as the most obvious CIN-tolerating mutation, it seems to enable both euploid and CIN cancers approximately evenly. This effect is likely a combination of the fact that cell-intrinsically, both CIN and Stat1^KO^ reduce growth of p53^KO^ cells, whereas *in vivo* Stat1^KO^ prevents efficient immune activation (but not infiltration). The detrimental effect of CIN we observed in our mouse experiments is reflected in a human breast cancer cohort, where patients with *TP53* mutant tumors show the overall worst survival without CIN, the opposite of *TP53* wild-type tumors. *TP53* mutant tumors, however, still show on average higher levels of aneuploidy, likely reflecting an imperfect correlation between aneuploidy and CIN in observational cohorts like the TCGA.

Myc activation, on the other hand, seems to be CIN-specific, as evidenced by the recurrent copy number amplifications both in our mouse cohort and across different human cancers. Strikingly, the lack of transposon insertions in Myc, combined with the lack of transcriptional upregulation of MYC in human cancers above its amplification status (*48*), suggests that CIN is required exactly because it enables these copy number amplifications and MYC as an oncogene is inert to epigenetic modulation. Previously, MYC-driven cancers have been described to be immune exclusionary by matter of the protein itself (*49*). Here, we find that with the pro-inflammatory effect of mis-segregations, cancer cells restrict Myc target transcription and will still be eliminated by both the innate (NK cells) and adaptive (T-cells) immune system. In order to overcome this, CIN-driven tumors need to dampen their interferon response, which they achieve through inactivation of Stat1 signaling. We showed that overexpression of the Myc oncogene and loss of Stat1 work synergistically in evading immune cells to sustain and promote the growth of CIN tumors in our mouse cohort and a clean engineered and allografted system in 3T3 cells, consistent with what we observe in human patient tumors.

We do not yet know how loss of STAT1 signaling is driven exactly in most cases, as mutations of *STAT1*, even in combination with *IFNA* loss, only account for a subset of tumors that were scored as STAT1-low by our gene expression signature. However, we can assume that alterations in other genes exist that mediate STAT1 inactivation, as evidenced by the striking insertion/mutation pattern in STAT1 interactors and the observed loss of inferred STAT1 activity. Indeed, it has been shown that for example Ras activation can counteract Myc-driven immunosurveillance (*50*). We believe this will be an important aspect of follow-up studies and a starting point to designing therapies targeted at cancer-intrinsic inflammation. This could be done for instance through reactivation of STAT1 or inhibition of specific interactors, likely in combination with immune checkpoint inhibitors (*51, 52*).

Finally, our data set provides a plethora of additional oncogenic hits that we did not explicitly follow up on, and we hope it will hence serve as a valuable resource for the CIN and aneuploidy community.

## Materials and Methods

### In vivo transposon screen

To generate euploid tumors, we made use of a PiggyBac (PB) transposon mouse strain (mixed C57BL/6 genetic background) (*53, 54*). Briefly, this strain harbors 15 transposable elements on chromosome 16 for which the transposon elements contain a promoter, splice donor (SD), bidirectional SV40 polyadenylation (PA) signals, and two splice acceptor (SA) signals flanked by PB inverted repeats, allowing for both loss and gain of function mutations depending on the integration site. To mobilize transposons in a tissue-specific manner, PB mice were crossed with mice expressing a conditionally activated transposase (Tpase), located at the Rosa26 locus on chr. 6 and preceded by a Lox-STOP-Lox sequence. In addition, these mice were crossed into an Mx1-Cre background so that PB Transposase (Tpase) can be activated specifically in the hematopoietic system following PolyI:PolyC treatment (*55*).

To generate mice that develop tumors with a CIN phenotype in the hematopoietic system, PB; Transposase; Mx1-Cre mice were crossed into a Mad2 conditional knockout (Mad2^f/f^) background, which promotes a strong CIN phenotype when switched by Cre-recombinase activity (*11, 22*). Next, we set up four cohorts of mice, one to yield euploid tumors (PB; Tpase; Mx1-Cre; 45 mice), one to yield tumors with a CIN phenotype (Mad2^f/f^; PB; Tpase; Mx1-Cre; 102 mice) and two control cohorts (Mad2^f/f^; Mx-Cre (10 mice) and Mad2^f/f^; PB; Tpase; Mx1-Cre (15 mice)). At the age of 8-12 weeks, the first three cohorts received 5 doses of 200 μg polyI:polyC (pI:pC; treatment every other day) by intraperitoneal injection to activate Mx1-Cre. The fourth cohort serves as a control and was left untreated.

We then monitored mice for signs of illness and euthanized mice by isoflurane sedation followed by cervical dislocation when they were at humane endpoints. Human endpoints included a larger than 20% weight loss within a period of two weeks, abnormal blood counts, palpable masses and dyspnea. Blood samples were collected at euthanasia via an orbital puncture into heparin-coated capillaries (Arstedt) and red blood cells removed by Erylysis (0.85% ammonium chloride) after which the pellet was stored at -80°C. Next, we inspected all organs with particular attention to the spleen, thymus, lymph nodes, liver and kidney. In addition to blood samples, we stored samples of the ear, tail, spleen, liver and bone marrow and any other macroscopically abnormal tissue. Tissues were then processed using standardized procedures described below.

To isolate bone marrow, bones were crushed using a mortar and single cell suspensions were derived using a 100 μm strainer. Red blood cells were removed using Erylysis and cells were frozen in 10% DMSO (Sigma) in FBS. Spleen, thymus and lymph node samples were processed into four fractions: 1) a tissue fragment for pathology (stored in 4% formaldehyde (Sigma) in PBS (Life)), 2) a fraction for nucleus, RNA and DNA isolation (stored as single cell suspensions in FBS with 10% DMSO, 2) a fraction for flow cytometry (stored as a single cell suspension in Cytofix/Cytoperm (BD Biosciences)), and 4) a backup sample, stored as a tissue fragment at -80°C.

All animal protocols were approved by the Central Committee for Animal experiments (CCD; permits AVD105002016465 and AVD105002016466) and UMCG Committee for Animal Care (IvD).

### Flow cytometry

For flow cytometry assessment, single cell suspensions were (thawed and) washed with 1x D-PBS. We excluded dead cells in samples originating from transplanted 3T3 tumors using Zombie Aqua labeling (423102, Biolegend) according to the manufacturer’s protocol. We then stained cells in 1x D-PBS containing 2% Fetal Bovine Serum (FBS, Life) and the relevant surface antibodies (**Table S1**) for 30 minutes at 4°C, after which we washed the cells. Next, we analyzed the stained samples with flow cytometry on a BD LSR-II flow cytometer (BD Biosciences). Data was analyzed using FlowJo Software (TreeStar).

### Shallow whole genome sequencing to quantify aneuploidy

To determine the average karyotype of individual tumors, we sorted 30 nuclei from each stored tumor by flow cytometry from samples frozen in 10% DMSO in FBS (mini-bulk sequencing). In addition, to determine intratumor heterogeneity as a readout of ongoing CIN, we also sorted 24 individual nuclei for three tumors for single cell whole genome sequencing. For both sequencing approaches, we next prepared the sorted nuclei for pre-amplification-free mini-bulk/single-cell whole genome sequencing using a Bravo liquid handler platform (Agilent Technologies) as previously described (*56*). In short, DNA was fragmented using micrococcal nuclease (0.5U or 1.25U for single cell or mini-bulk resp.) followed by end-repair, A-tailing and Illumina adapter ligation. After AMPure XP bead clean-up, the adapter-containing DNA fragments were subjected to PCR amplification for 12 or 17 cycles (mini-bulk or single cells, respectively) using custom multiplexing PCR primers to incorporate unique library-specific barcodes. Pooled libraries were sequenced on a NextSeq 500 system (Illumina; 77 cycles; single-end).

Data analysis was performed with the *AneuFinder* software package (*57*) version 1.18. For single-cell WGS, we used inferred copy numbers and karyotype measures (aneuploidy, heterogeneity) from AneuFinder directly. For mini-bulk WGS, we assumed that the copy number of the median segment is euploid, and quantified aneuploidy as the average absolute deviation from that copy number (up to a score of 0.2, at which point we considered a sample fully aneuploid).

### Transposon insertion sequencing

To determine transposon integration sites, we isolated DNA from single cell suspensions or single cell pellets using a DNeasy Kit (Qiagen) according to manufacturer’s protocol. To detect transposon integration sites, we used QISeq (*2*). In short, genomic DNA was isolated, sheared to a mean fragment length of 250-bp on a Covaris AFA sonicator and ligated to a custom Splinkerette adapter. Transposon-containing fragments were enriched with 12 cycles of transposon-specific PCR for both the 5’ and the 3’ transposon ends in separate libraries. Bar coding of individual samples and completion of Illumina adaptor sequences was achieved with an additional 8 cycles of transposon-specific PCR and a custom array of 96-unique bar-coding primers. After size selection via magnetic bead purification (Beckman Ampure XP) and quantification with qPCR, libraries were pooled equimolarly in two separate 5’ and 3’ pools and sequenced on a MiSeq Desktop Sequencer (Illumina).

### Mapping Common Insertion Sites (CIS) and aneuploidy-specific CIS

We downloaded mouse gene annotations for GRCm38 from Ensembl 102, and subset to all protein coding genes on chromosomes 1 to 19 and X. We defined as gene insertions any PiggyBac insertion in the gene body (defined by start and end position of the gene) and up to 10 kb upstream, excluding the genes *Sfi1*, *Drg1* and *Eif4enif1* (as part of a region with a high number of false positives). For DNA insertions, we counted the number of TTAA target sites for each gene and the genome as a whole (on the selected chromosomes). We defined a genome-wide base pair background insertion rate by the total number of transposon insertions, and tested the number of insertions in each gene compared to this rate using a Poisson test. Genes where we observed a significant enrichment over the background rate were called Common Insertion Sites (CIS). Additionally, we quantified the presence of RNA evidence for any insertion using *STAR-fusion* 2.7.9a (*58*) and the *IMfusion* tool (*59*) using a combined genome of GRCm38 (Ensembl 92) and the transposon sequence of pA6-GrOnc (*34*). For aneuploidy-specific CIS, we tested whether the number of insertions scales with aneuploidy (at a maximum value of 0.2, at which point samples re considered fully aneuploid) using a linear model for all genes as before.

To narrow down the list of targets for a potential follow-up, we used all human interactions from OmniPath (*60*) and got mouse orthologs using Ensembl 102 as recommended in the documentation. We converted this graph to a simple undirected network (only one possible edge between nodes, no self-edges). We then used the *BioNet* package (version 1.50.0) (*61*) to extract the main interacting component of all CIS or aneuploidy-specific CIS using the formula where the score for each node *S* = −*log*10(*S*) + *log*10(*t*). Here, *s* refers to the FDR-adjusted p-value of each CIS or unadjusted p-value for aneuploidy specific CIS. *t* refers to the threshold at which scores are considered positive and is set to 0.1and 0.05 for CIS and aneuploidy-specific CIS, respectively. In the extracted subnetwork, we calculated the Hub centrality of each node using the *igraph* package (version 1.2.6). For visualizing the combined core network between CIS and aneuploidy-specific CIS, we combined both networks and removed nodes with a degree of 2 or lower, followed by a second iteration of genes with a degree of 1 or lower.

### RNA isolation, RT PCR, RNA sequencing and analysis

We isolated RNA using the RNAeasy plus mini kit (Qiagen, 74136) according to manufacturer’s protocol. RNA was used for quantitative reverse transcription-PCR (RT-PCR) and RNA-sequencing. For RT-PCR, we synthesized complementary DNA (cDNA) using a mixture of M-MuLV reverse transcriptase (NEB, M0253L), RNAse inhibitor (NEB, M0307L) and random primers (NEB, S1330S) according to the manufacturer’s protocol. We performed RT-PCR in technical triplicates using the iTaq Universal SYBR Green Supermix (Biorad, 1725150) and used primers listed in **Table S2**. As reference genes, we used Tubulin or Actin. RT-PCR experiments were performed on a LightCycler® 480 Instrument (Roche).

For RNA-sequencing, we first examined RNA quality using the Agilent 2100 Bioanalyzer (Agilent Technologies). We typically used 300 ng of total RNA as input for enrichment by NEXTflex Poly(A) Beads (Bioo Scientific) followed by library preparation using NEXTflex Rapid Directional qRNA-Seq Kit (Bioo Scientific). Up to 32 libraries were pooled and sequenced to 450 million reads on an NextSeq sequencer (Illumina).

We aligned FASTQ files to Ensembl 92 GTF for GRCm38 using STAR 2.7.9a (*8*) and counted the number of reads for each gene using *subread* 1.6.1 (*62*). We used the *DESeq2* 1.31.3 (*63*) variance stabilizing transformation as a basis for Principal Component Analysis (PCA) and UMAP dimensionality reduction using the *uwot* package (version 0.1.10). We calculated differential expression of all samples with aneuploidy using a Wald test aneuploidy score, correcting for the cancer type. For differential expression of gene sets, we used a linear model to quantify whether genes in a set have a different Wald statistic compared to genes not in the set. As gene sets we used the *MSigDB Hallmark 2020* collection from *Enrichr* (*64*) and *DoRothEA* transcription factor-target relationships (*65*). To obtain sample-level scores for individual gene sets, we used *Gene Set Variation Analysis* (*66*). We performed all conditional tests using these GSVA scores and a linear model.

For the comparison of our mouse tumors with human leukemias, we used the *Microarray Studies in Leukemia* (MILE) database (*38*), obtained via *ArrayExpress* (*67*) and processed via the *oligo* package (*68*) version 1.54.1.

### Cell culture

BT549 and MCF7 cell lines were cultured in RPMI 1640 medium with GlutaMAX supplement (Life), 10%FBS, and 100 U/ml penicillin/streptomycin (P/S). NIH/3T3 cell lines were cultured in DMEM (Invitrogen), 10% FBS, MEM non-essential amino acids solution, β-Mercapethanol (β-ME), and 100 U/ml penicillin/streptomycin (P/S). NK92 cell lines were cultured in RPMI 1640 medium with Glutamax, 12.5% FBS, 12.5 % Horse Serum, 100 U/ml human interleukin 2 (Peprotech, 200-02), and 100 U/ml P/S. All cells were grown at 37°C in the presence of 5% CO_2_ in a humidified environment.

### Genetic modifications using CRISPR/Cas9

To engineer Stat1/STAT1 and p53 KO cells, we designed CRISPR guideRNAs targeting exon 5 of STAT1 or Stat1 (**Table S2**) and exon 4 of p53 (**Table S2**). We cloned guideRNAs into the pSpCas9(BB)-2A-Puro V2.0 (PX459) plasmid using BbsI restriction sites (Addgene plasmid #62988, (*69*)). To generate STAT1/Stat1 and p53 knockout BT549, MCF7, and NIH/3T3 cells, cells were transfected with 2 μg of the respective guideRNA plasmids using FuGene (Promega) according to the manufacturer’s protocol. 48 hours after transfection, cells were selected with puromycin (InVivogen; 1 μg/ml for BT549 and MCF7 and 2 μg/ml for NIH/3T3) for 2 days before validating the knockouts by immunoblotting.

### Cloning of MYC overexpression plasmid

To generate a constitutive MYC overexpression plasmid, MYC was amplified from a human cDNA library generated from RPE1 cells by PCR (primers listed in **Table S2**) and cloned into a pMCSV plasmid using BamHI and EcoRI restriction sites. pBCH-KIF2C and pBCH-dnMCAK were generated by transferring KIF2C and dnMCAK fragments isolated from expression plasmids (*41*) into a pBCH lentiviral backbone using NdeI and BamHI.

### Protein quantification using immunoblotting

To determine whether cells indeed had lost protein expression of the CRISPR-targeted genes (STAT1/Stat1; p53), we performed Western blots. For this, we lysed equal numbers of cells using ELB lysis buffer (150 mM NaCl, 50 mM Hepes pH 7.5, 5 mM EDTA, 0.1% NP-40) supplemented with protease (Roche, 11697498001) and phosphatase inhibitor (Sigma Aldrich, 4906845001). We then mixed the cell lysates with a 5X sample buffer (50% Glycerol, 10% SDS, 05M DTT, 250 mM Tris pH 6.8) and boiled samples for 5 minutes at 98°C. Samples were run on SDS-polyacrylamide gels (SDS-PAGE) and then transferred onto a polyvinylidene difluoride (PVDF) membrane (Millipore). We blocked protein-containing membranes with Odyssey blocking buffer (LiCor) diluted 1:1 with TBS for 30 minutes at room temperature. Blots were incubated overnight at 4°C in blocking buffer containing the primary antibodies (**Table S3**). The next day, we washed membranes for 3 times (5 minutes each) with TBS-T (19 mM Tris base, NaCl 137 mM, KCl 2.7 mM and 0.1% Tween-20) and incubated them with fluorophore-conjugated secondary antibodies: IRDye 800CW Goat anti-Rabbit IgG (H + L) (Licor, 1:15.000) and IRDye 680RD Goat anti-Mouse IgG (H + L) (Licor, 1:20.000) for 1 hour at room temperature. Membranes were washed for 3 times (5 minutes each) with TBS-T followed by membrane quantification using an Odyssey CLx fluorescence scanner (Licor). Finally, we quantified protein expression using ImageStudio Lite software (Licor).

### Retroviral and lentiviral transductions

To produce retrovirus, we transfected Phoenix Ampho cells (ATCC CRL-3213) with 3 μg of plasmid (pMCSV c-Myc or empty vector). To produce lentivirus, 293T cells were transfected with lentiviral packaging vectors (3 μg pSPAX2 (Addgene plasmid, 12260) and 1 μg pMD2.G (Addgene plasmid, 12259) combined with the overexpression vectors H2B-cherry, pBCH KIF2C or pbCH dnMCAK. 48 hours post-transfection, we filtered medium from transfected 293T/Phoenix Ampho cells through a 0.45 μm filter (Corning, 431220) and transferred medium to the target cells. To increase transduction efficiency, 8 μg/ml polybrene (Sigma) was added.

### Proliferation assays

To quantify proliferation following mock- or reversine treatment, 20,000 cells BT549 (WT or STAT1^KO^) or MCF7 (WT, p53^KO^, STAT1^KO^, or p53 STAT1^KO^) were seeded 12 well plates. 24 hours later, the medium was supplemented with 500 nM Reversine (Sigma Aldrich, R3904) or DMSO (AppliChem, 67685) for 72 hours. Next, cells were trypsinized and counted using a TC20 automated cell counter (Biorad, 145-0101). We then calculated cell proliferation by normalizing cell counts to the DMSO control using Microsoft Excel.

For 3T3 cells, 2,000 3T3 cells (genotypes as indicated) were seeded in a 6-well plate. Medium was replenished every 3 days and after 10 days of incubation, cells were washed two times with ice cold 1xD-PBS. We then fixed cells using 4% formaldehyde in PBS and stained cells with 0.1% crystal violet (Sigma Aldrich, C3886) diluted in water. Growth dynamics were quantified by dissolving crystal violet in 10% acetic acid followed by quantification of the absorbance at 590 nm. Relative proliferation rates were then calculated using Microsoft Excel.

### Co-culture assays

For NK92 co-culture assays, 10,000 cells BT549 (WT or STAT1^KO^) or MCF7 (WT, p53^KO^, STAT1^KO^, or p53;STAT1^DKO^) were seeded in 24 well plates. After 24 hours, we added either 250 nM reversine or DMSO. 48 hours later, NK92 cells were added to the mock- or Reversine-treated cells at a ratio 1:2 (NK92:BT549) and co-cultured for 48 hours. Then, we removed NK92 cells by washing out the wells with PBS twice, and harvested and counted the remaining BT549 and MCF7 cells using a TC20 automated cell counter (BioRad). The effect size of the NK92 cell co-culture was calculated by normalizing cell numbers to BT549/MCF7 mock or reversine-treated cultures that were not co-cultured with NK92 cells.

For human Peripheral Blood Mononuclear Cells (PBMC) transwell assays, we seeded BT549 (WT or STAT1^KO^) in a 24-well plate (10,000 cells per well) and treated cells with DMSO or reversine for 96 hours. We isolated PBMCs from Buffycoat derived from healthy volunteers using the Ficoll-Paque method (*70*). For the transwell assay, we next seeded 2×10^6^ human PBMCs on top of a filter membrane of a transwell insert (6.5 mm Transwell with 3.0 μm pore, Corning). After a 24- or 48-hours incubation period, we harvested the medium from the bottom compartment and counted the number of migrated cells using a Bürker-Türk counting chamber (Merck, BR719505). After cell counting, the migrated PBMCs were collected by centrifugation and stained for flow cytometry as described above (see flow cytometry section).

For mouse 3T3 transwell assays, we seeded NIH/3T3 cells of indicated genotype (see Figure 4) into 24-well plates (10,000 cells per well). Mouse splenocytes were isolated from wild type Balb/c mice (7-10 weeks of age). For this, we homogenized mouse spleen and passed the homogenate through a 70 µm Cell Strainer (Corning, 431751). We next centrifuged the resulting cell suspension at 300g, followed by incubation with 1x red blood cell lysis buffer (Biolegend, 420301) for 5 minutes on ice. After this, splenocytes were centrifuged and the pellet was resuspended in RPMI medium. Next, 2×10^6^ splenocytes were seeded on transwell as described for human PBMC co-cultures. Splenocyte migration was quantified after a 48 hours incubation period as described for human PBMC co-culture assays and migrated splenocytes were processed for flow cytometry staining as described above.

### Time-lapse microscopy imaging

To quantify chromosome missegregation rates in cultured cells, we seeded 50,000 NIH/3T3, MCF7 or BT549 cells with or without genetic modifications and transduced with an H2B-mCherry construct into a 4-section imaging chamber (Greiner Bio-one,627870). Cells were grown overnight and treated with mock-control or 250 nM reversine 1 hour before imaging started. We imaged cells for at least 16 hours using a DeltaVision Elite (GE Healthcare) microscope equipped with a CoolSNAP HQ2 camera and a 20x 0.75 NA or 40x 0.6 NA immersion objective. Images were acquired every 7 minutes, including z-stacks of 20 images at 0.4 μm intervals. Image analysis was done using ICY software (Institut Pasteur) and included all cells that entered mitosis and stayed in frame throughout the imaging session.

### 3T3 inoculation and tumor transplantation

Tumor grafting experiments were performed in 7 to 11 weeks old nude athymic or Balb/c mice and 3T3 tumors were pre-grown in immunodeficient mice. For this, recipient nude (Hsd:Athymic Nude-Foxn1 <NU>) mice were anesthetized using isofluorane following which we injected 0.5×10^6^ NIH/3T3 cells of the following genotypes: Myc^OEX^;KIF2C, Myc^OEX^;dnMCAK, Myc^OEX^;Stat1^KO^;KIF2C, Myc^OEX^;Stat1^KO^; dnMCAK, p53^KO^;KIF2C, p53^KO^;dnMCAK, p53^KO^;Stat1^KO^;KIF2C, and p53^KO^;Stat1^KO^;dnMCAK subcutaneously into the flanks of the mice. We carefully followed mice for tumor development and euthanized them when tumors reached 0.8 cm^3^ in diameter to harvest the primary tumors. Following tumor dissection in a petri dish containing RPMI medium, we chopped the isolated tumors into fragments of approximately 50 mm^3^. Next, we transplanted single 50 mm^3^ tumor fragments subcutaneously into the flanks of wild-type Balb/c mice. For this, we anesthetized recipient mice using isoflurane and placed them on a heating pad to maintain body temperature during surgery. During surgery, mice received 5 mg/kg of carprofen to minimize pain, and eye ointment to prevent dehydration of the conjunctiva. To prepare for tumor fragment transplantation, we shaved the skin of the neck, disinfected the surgery area and made a 1 cm skin incision. We created a subcutaneous pocket in each side of the lower back using a forceps, in which we deposited one tumor fragment each. Finally, we closed the skin with absorbable sutures. We carefully measured tumor growth every other day using a digital caliper. Tumor volume was calculated as *volume* = (*length* * *width*^2^)/2. Mice were euthanized at humane endpoints, *i.e.* when tumors would reach a size of 0.8 cm^3^. Following tumor dissection, we prepared single cell suspensions by homogenizing tumors through a 70 μm strainer. Cells were then washed in 1x D-PBS and stored as a single cell suspension.

### Histological assessment of 3T3 tumors

For histological assessment, we fixed tumor fragments in 4% paraformaldehyde (PFA; Sigma) overnight at 4°C. Subsequently, we dehydrated tissues in ethanol, processed and embedded them in paraffin according to standard histology procedures. To prepare samples for immunohistochemistry, we sliced 5 μm sections of tumors on a microtome. We dewaxed sections in xylene, and next cleared them in ethanol and rehydrated them in water followed by antigen retrieval in 10 mM citrate at a pH of 6.0 in a microwave for 15 minutes at 650 Watt. Slides were cooled down for 30 minutes at room temperature and washed with 2x with 1x D-PBS. Endogenous peroxidase was blocked in 1% H_2_O_2_ (Sigma) in 1x D-PBS for 30 minutes at room temperature in the dark. Next, we washed sections, and blocked them with normal goat serum in 1x D-PBS for 30 minutes, followed by overnight staining at 4°C with rat anti-mouse CD45 primary antibody (103102, Biolegend) diluted 1:500 in 1x D-PBS. The following day, slides were washed 3 times with 1x D-PBS containing 0.1% Tween 20 and incubated with goat anti-rat biotinylated secondary antibody diluted 1:125 in 1x D-PBS for 30 minutes at room temperature. After washing 3 times with 1x D-PBS containing 1% Tween 20, we incubated sections with ABC-peroxidase solution for 30 minutes at room temperature. Slides were then washed 3 times with PBS, cleared with water and incubated 10 minutes in the dark with 7 mM DAB 0.03% H_2_O_2_ in Tris HCl at a pH of 7.8 for 10 minutes at room temperature protected from light. After washing with tap water, we counterstained slides with hematoxylin for 30 seconds. Finally, slides were washed with tap water, dehydrated with ethanol, cleared with xylene and mounted using toluene mounting media. Whole slide imaging was performed using a NanoZoomer S60 Digital slide scanner (Hamamatsu) and analyzed with NDP-view2 software (Hamamatsu). Blinded manual counting of CD45^+^ cells was done using NDP-view2 software.

### TCGA and BT549 analysis

We obtained RNA-seq reads, Single Nucleotide Variants (SNVs, “mutations”), segmental copy number, and clinical data from the TCGA cohorts via the *TCGAbiolinks* package (*71*) 2.18.0. We quantified the degree of aneuploidy as the average absolute deviation from the euploid. We obtained sample purity estimates for these samples from the *ESTIMATE* publication (*72*) and scored gene set expression as described for the transposon cohort. We defined 10-year overall survival as the maximum duration to either a patient’s death or their last follow-up date, where we considered all patients as “alive” at 10 years should this date be longer than 10 years.

We calculated the change of Myc targets V1 (*MSigDB Hallmarks*) and STAT1 target expression (*DoRothEA*) with aneuploidy using a linear model between the two and correcting for sample purity for each cohort, each cohort split by p53 status, and all cohorts using the cohort as a covariate. For all associations, we only included primary tumors (barcode portion 01A), to exclude confounding by metastatic or non-tumor samples. We show the standard error for these estimates as bars for both gene sets, for the reader to gain an understanding of their confidence.

We quantified tumor samples as having STAT1 mutations if they are associated with the *STAT1* gene symbol in the variant catalogue and are listed as non-silent mutations. For obtaining interactors, we considered each tumor sample to have a STAT1 interactor mutation if there is a non-silent mutation in any of the first-degree neighbors as defined by *OmniPath* (*60*). We quantified the difference in euploid and aneuploid samples by applying a cutoff to the aneuploidy score of 0.1. We tested for the difference of *STAT1* mutations as a fraction of total mutations using a Wilcoxon Rank Sum Test. For testing for a difference between interactor mutations, we used the *mgcv* package (*73*) version 1.8-37 to fit a smooth Generalized Additive Model (GAM) to the relation between the number of total mutations and the number of STAT1 interactor mutations, including a fixed term for whether this sample belongs to a euploid or an aneuploid tumor. We obtained the significance for this fixed effect using an F test.

For our parental and genetically modified BT549 cell lines, we obtained gene counts as described for the transposon mouse cohort, except that we mapped to GRCh38 and additionally scored our gene expression using Gene Ontology 2021 gene sets from *Enrichr* (*64*). We compared the Wald statistic (the differential gene set expression estimate divided by its standard error) to the one we obtained from the gene set differential expression of our transposon cohort and quantified the concordance or discordance using a Fisher’s Exact Test on gene sets that are changing in both conditions (absolute Wald statistic > 1.5; colored points in **Fig. 5d-f**). We then obtained a “Acute CIN” or ”Inflammatory CIN” gene expression signature (top 70 genes BT549 Reversine vs. DMSO) and a “STAT1 KO CIN” gene expression signature (top 70 genes Reversine-treated BT549 STAT1^KO^ vs. STAT1^WT^), which we used to score all primary TCGA breast tumor (BRCA cohort) samples using GSVA. We confirmed that our “STAT1 KO” signature scores correlated significantly with scores obtained from previously published CIN signatures (CIN70, HET70) and aneuploidy. We then divided the cohort into “CIN” (acute CIN signature score is positive), “no CIN” (acute CIN signature is negative and STAT1 KO signature is negative), and “STAT1 KO CIN” (acute CIN signature is negative and STAT1 KO signature is positive), and compared patient survival between these groups in either the p53 wild-type or the p53 mutant setting. We quantified the survival differences using a Cox Proportional Hazards model, regressing out age at diagnosis and sample purity. In addition, we compared the associations to what we could obtain when replacing the “STAT1 KO CIN” signature with previously published CIN signatures (CIN70, HET70), aneuploidy, or a proliferation (E2F Targets Hallmark) signature.

Finally, we quantified how well different factors are able to predict MYC expression and MYC target expression in the TCGA BRCA cohort. For this, we chose the following predictors: *MYC* copy number, *TP53* mutation status, “STAT1 KO CIN” signature score, PAM50 subtype, and MYC gene expression (for targets only). We tested each predictor individually for the whole cohort and split by *TP53* mutation status using a linear model. We also tested the contribution of each individual predictor taking into account all others (Type II ANOVA applied on the multivariate regression models, *car* package 3.0-11).

## Supplementary Tables

**Table S1.** Flow cytometry antibodies

| Antibody | Clone number | Distributor |
| --- | --- | --- |
| CD45.2-PE | 104 | Biolegend |
| CD4-FITC | GK1.5 | Biolegend |
| CD8a-PerCPCy5.5 | 53-6.7 | Biolegend |
| CD19-APC | 1D3/CD19 | Biolegend |
| CD335(Nkp46)-APC Cy7 | 29A1.4 | Biolegend |
| CD45-PerCPCy5.5 | 30-F11 | Biolegend |
| CD3-APC | UCHT1 | Biolegend |
| B220-AF700 | RA3-6B2 | Biolegend |
| Gr1-PECy7 | RB6-8C5 | Biolegend |
| Mac-1-PECy7 | M1-70 | Biolegend |
| ckit-PE | 2B8 | Biolegend |
| Sca-1-BV421 | D7 | Biolegend |

**Table S2.**
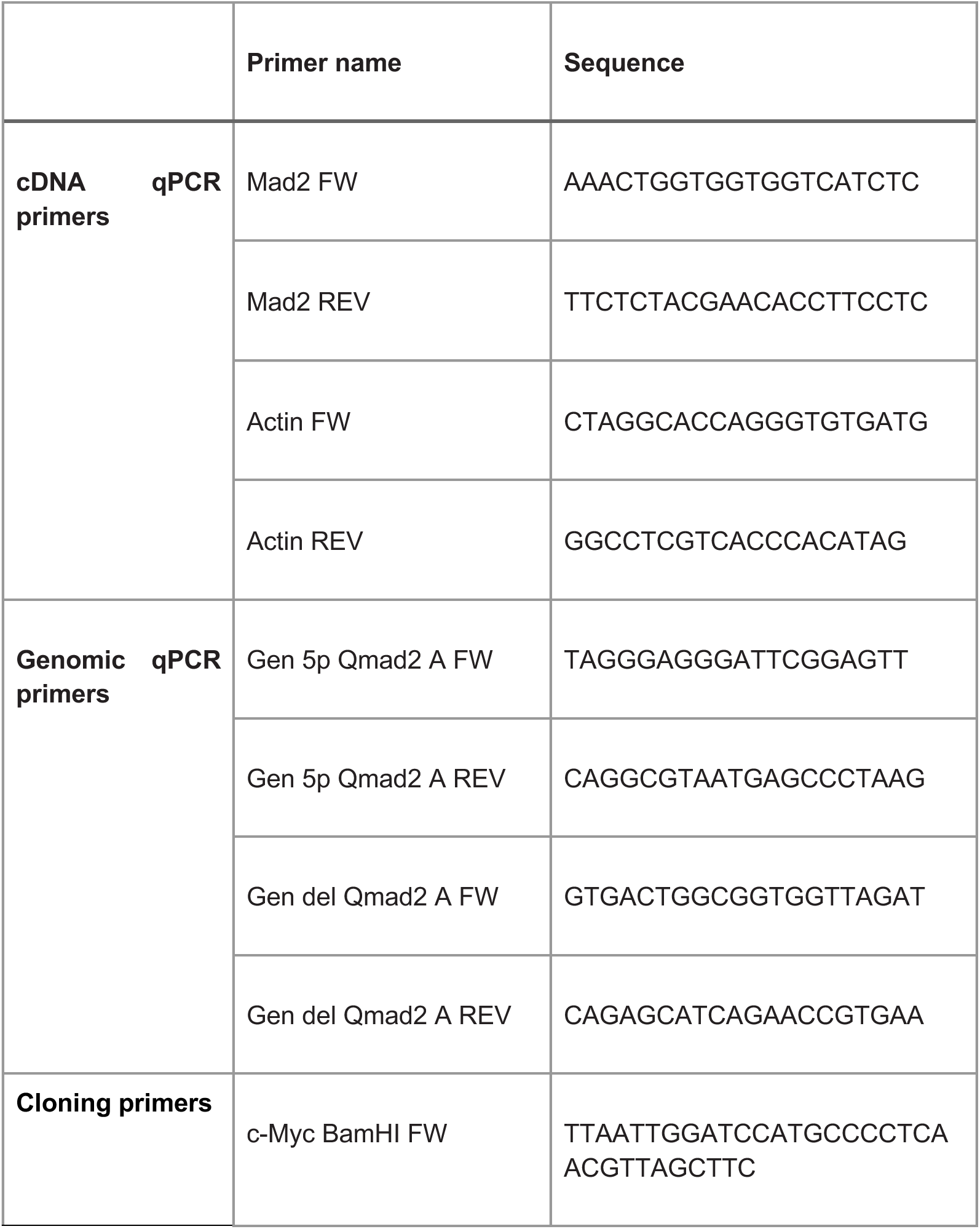
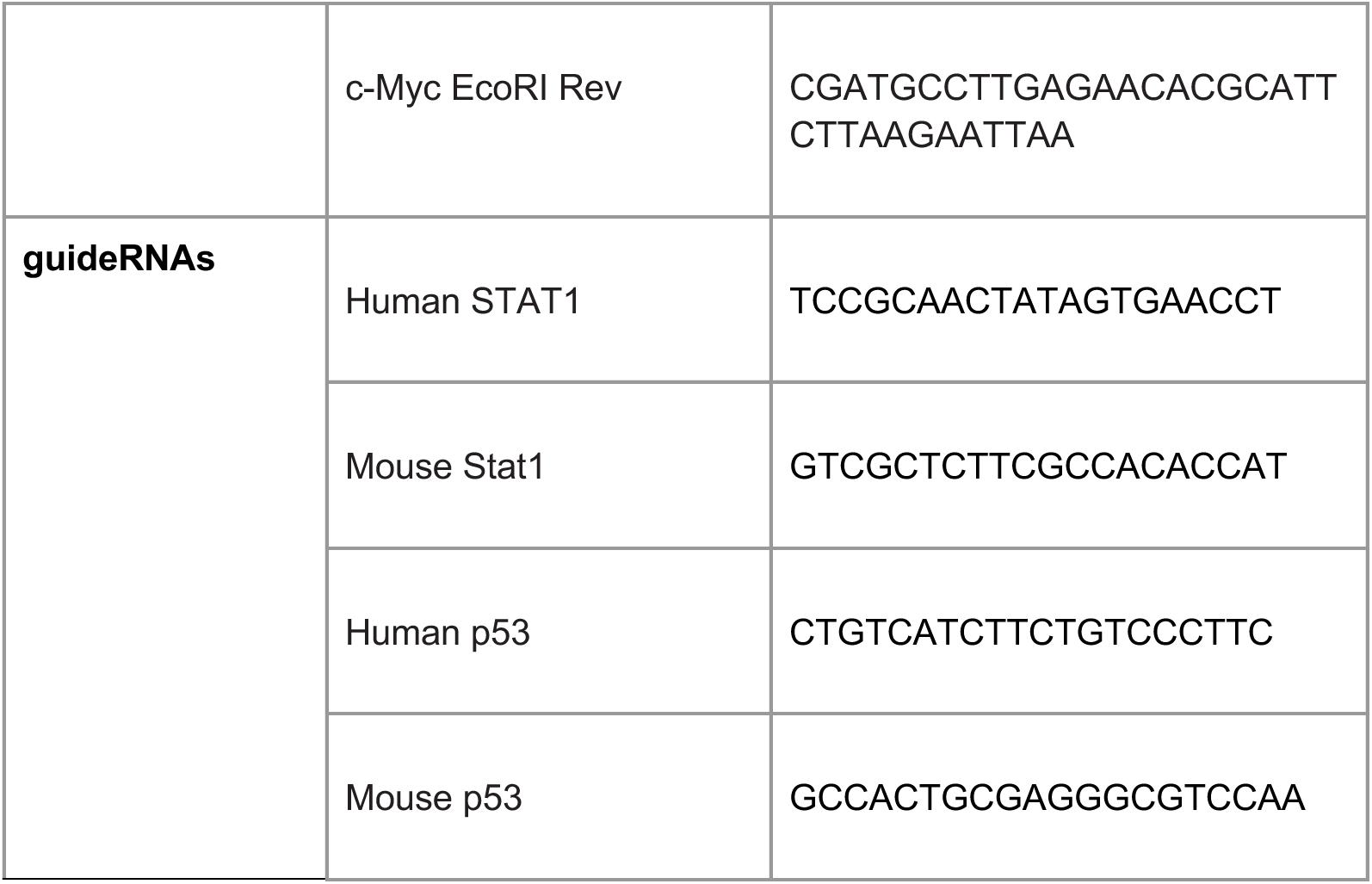
Primer and guide sequences

**Table S3.** Immunoblotting antibodies

| **Antibody** | **Clone number** | **Distributor** |
| --- | --- | --- |
| Mad2 | 61078 | BD Bioscience |
| STAT1 | 9175 | Cell Signaling |
| p53 | sc-126 | Santa Cruz |
| c-Myc | 5605 | Cell Signaling |
| β-actin | 4970 | Cell Signaling |

## Acknowledgements

We thank Michel Weij and Daryll Eichorn at the central animal facility for assistance with animal experiments and Johan Teunis and Theo Bijma at the flow cytometry facility for help with flow cytometry. We thank Rianna Arjaans and Nancy Halsema at the UMCG/ERIBA Research Sequencing facility for help with RNA sequencing library preparation and sequencing and Mathilde Broekhuis and Jonas Seiler at the UMCG/ERIBA iPSC/CRISPR facility for advice on the CRISPR KO experiments. We are grateful to Jiske Tiersma, Johanna Veldmann, Arthur Svendsen, Bertien Dethmers-Ausema, Maurits Roorda, Francien Talens and Chloe Lee for general advice on the project. We thank the members of the Foijer lab for fruitful discussions.

## Author contributions

Conceptualization: M.S., C.H., L.J.J., J.E.S., F.F.

Experimental design and methodology: M.S., C.H., L.J.J., M.R.R, F.F.

Computational analyses and data integration: M.S.

Mutagenesis screen and tumor harvest L.J.J., J.E.S., B.B, P.L.B.

Mouse pathology: M.H.K., A.d.B.

Integration site sequencing: J.L.C, A.D., A.S. J.E.S.

Mapping common insertion sites M.S., M.B., T.E. and H.P.

*In vitro* experiments C.H., L.J.J., A.E.T, P.L.B, A.v.d.B, L.A.R, S.G., D.L.

RNA, mini-bulk and scWGS sequencing: D.C.J.S.

*In vivo* validation experiments: C.H. and M.R.R;

Supervision: H.W.N, M.d.B., G.d.H. R.H.M., M.C.T, R.R., J.L.C., F.F.

Provided reagents: G.d.H., R.R., J.L.C., M.A.T.M.v.V.

Funding acquisition: F.F.

Writing original draft: M.S., C.H., F.F.

Writing edits: M.R.R., M.A.T.M.v.V, M.C.T., G.C.V

## Supplementary Figure Legends

**Figure S1.** Basic characterization of tumors arising in transposon mutagenesis screen mice. **(a)** Representative H&E stainings for the three main types of identified malignancies: acute T-cell lymphoma (T-ALL, left panel), B-cell tumors (middle panel) and myeloid tumors (histiocytic sarcoma, right panel). **(b-d)** Flow cytometry gating strategy for T-ALLs **(b)**, B-cell tumors **(c)** and myeloid tumors **(d)** showing representative examples for each. **(e)** Genomic PCR to quantify Mad2 deletion products in a selection of the Mad2 transposon malignancies. **(f)** Mad2 switching in sporadic tumors arising in uninduced transposon tumors. **(g)** Quantitative genomic PCR showing Mad2 deletion in Mad2 transposon tumors. **(h)** Quantitative RT-PCR showing loss of Mad2 RNA expression in Mad2 transposon tumors. **(i)** Western blot showing loss of Mad2 protein in a selection of Mad2 transposon tumors, including 2 tumors that did not show Mad2 protein loss.

**Figure S2.** Transposon insertion sites. **(a)** Volcano plot of Common Insertions Sites (CIS) with log2 fold change in a given gene *vs.* the genome background. **(b)** Volcano plot of aneuploidy-specific CIS by testing for a difference in aneuploidy level in tumors that have an insertion *vs.* tumors that do not. Points with a black border are significant in the other respective association. **(c)** Protein-protein interaction subnetwork for all CIS with highest sample numbers and hub centrality (right) annotated. **(d)** Subnetwork of aneuploidy-specific CIS, additionally showing bias towards aneuploid (blue) or euploid tumors (red). Dark grey bars indicate that these associations are also present in the other respective network.

**Figure S3.** RNA sequencing of tumors. **(a)** Principal Component Analysis (PCA) of mouse tumors, sequentially resolving marker sets for T-ALLs, Myeloid leukemias, and subtypes of B-like ALLs driven by Ebf1, Ets1, or Erg. **(b)** Ets1 and Erg expression in the transposon cohort compared to a large human leukemia cohort (MILE) shows a consistent pattern between T-ALLs, Myeloid leukemias, and B-like precursor ALLs/hyperdiploid ALLs. **(c)** Pattern of aneuploidies across human leukemias and our mouse tumors shows that mature B-ALLs are euploid in the human cohort, while B precursor ALLs show a range of different aneuploidy levels, consistent with our mouse cohort. Myeloid leukemias have many more euploid samples in both cohorts. T-ALLs can be aneuploid in both cohorts, but tend to be more aneuploid in our mouse tumors.

**Figure S4.** Characterization of engineered 3T3 cell lines. **(a)** 3T3 Stat1^KO^ cells were generated using a guideRNA targeting the Stat1 gene. Cell lysates from wild type or Stat1^KO^ 3T3 cells were immunoblotted to verify gene inactivation. **(b)** 3T3 cells were retrovirally transduced with a Myc overexpression plasmid to generate Myc^OEX^ lines. Cell lysates from wild type or Myc^OEX^ 3T3 cells were immunoblotted for Myc expression to confirm overexpression. **(c)** A guideRNA targeting the p53 gene was transfected into 3T3 cells to obtain a 3T3 p53^KO^ line. Gene inactivation was confirmed by a Western blot. **(d-f)** Live cell imaging of **(d)** wild type, **(e)** Myc overexpressing or **(f)** p53^KO^ 3T3 cells expressing KIF2C or dnMCAK confirming CIN^low^ or CIN^high^ phenotypes, respectively.

**Figure S5.** Stat1 inactivation promotes outgrowth of tumors with a CIN phenotype by suppressing CIN-induced immune infiltration in a Myc^OEX^ background and by preventing CIN-induced immune cell activation in Myc^OEX^ and p53^KO^ backgrounds. **(a-b)** Tumor growth of Myc^OEX^ (left panel) or p53^KO^ (right panel) tumor fragment allografts into immunocompetent Balb/c mice. CIN^high^ tumors display a growth delay compared to CIN^low^ tumors in a Stat1 proficient background, which is alleviated in a Stat1 deficient background. **(c)** End mass of tumors isolated from allografted Balb/c mice derived from Myc^OEX^ (left panel) or p53^KO^ 3T3 cell lines (right panel). **(d)** Immunohistochemistry to detect CD45^+^ cells infiltrated into the allografted tumors revealed similar infiltration rates as determined by flow cytometry (**Fig. 4** **i,j**). **(e)** Quantification of the fraction of CD45^+^ cells as determined by immunohistochemistry for all genotypes (left panel, Myc^OEX^ and right panel p53^KO^). For each tumor, two to four areas were quantified. **(f-i)** Quantification of **(f)** CD4^+^ T-cells, **(g)** CD8^+^ T-cells, **(h)** NK cells, or **(i)** CD19^+^ B-cells observed in Myc^OEX^ (left panel) or p53^KO^ tumors (right panel) as determined by flow cytometry. Tumors were stratified according to size, as growing (> 200 mm^3^ or non-growing (<200 mm^3^). Small Myc^OEX^ tumors that display CIN show increased fractions of CD4^+^ and CD8^+^ cells, while p53^KO^ tumors with CIN show an increased fraction of CD4^+^ and CD19^+^ cells. Bars in **(c,e-i)** represent mean +/- s.e.m, **(c,f-i)** significance was tested by two-sided t-tests.

**Figure S6.** STAT1 loss dampens CIN-induced inflammatory signaling, immune cell recruitment, and NK92 cytotoxicity in human breast cancer cells. **(a)** Live cell imaging of BT549 cells shows that CIN induced by 250 or 500 nM reversine increases the rate of mitotic abnormalities. **(b)** BT549 STAT1^KO^ cells were generated using a guideRNA targeting the human STAT1 gene. Immunoblots against STAT1 protein were performed to confirm gene inactivation. **(c)** Comparing the proliferation rates of wild type and STAT1^KO^ BT549 cells shows no significant growth difference. **(d)** Live cell imaging of wild type or STAT1^KO^ BT549 cells shows no difference in chromosome mis-segregation rates. **(e)** Proliferation rates of STAT1^KO^ cells are less affected than wild type BT549 cells by acute CIN provoked by 500 nM reversine treatment. **(f)** STAT1^KO^ cells show less apoptosis compared to wild type BT549 cells, quantified by Annexin V staining and flow cytometry. **(g)** Transwell migration assays reveal that STAT1^KO^ BT549 cells attract fewer immune cells than wild type BT549 cells following acute CIN induced by 250 nM reversine. **(h)** STAT1^KO^ cells are killed less than wild type BT540 cells by cytotoxic NK92 cells following acute CIN induced by 250 nM Reversine. Co-culture ratio was 1:2 (NK92: BT549). **(i)** Live cell imaging shows that acute CIN induced by 250 or 500 nM reversine induces mitotic abnormalities in MCF7 cells. **(j)** MCF7 cells were transfected with a guideRNA targeting the STAT1 gene to generate MCF7 STAT1^KO^ cells. Immunoblots confirm loss of STAT1 expression. **(k)** Comparing growth rates of wild type, p53^KO^, STAT1^KO^, or p53;STAT1^DKO^ MCF7 cells demonstrates that p53 inactivation promotes proliferation. **(l)** p53^KO^ or/and STAT1^KO^ cells show moderately increased tolerance towards acute CIN induced by 500 nM reversine. **(m)** STAT1^KO^ and STAT1;p53^DKO^ MCF7 cells are killed less efficiently by cytotoxic NK92 cells than wild type and p53^KO^ MCF7 cells following acute CIN induced by 250 nM reversine. Co-culture ratio was 1:1 (MCF7 cells: NK92 cells). Bars in (**c,e-h, k-m**) represent mean +/- s.e.m, significance was tested by two-sided t-tests.

**Figure S7.** TCGA breast cancer analysis. **(a)** The BT549-derived STAT1 knockout signature significantly correlates with aneuploidy and previously published CIN signatures such as CIN70 and HET70. **(b)** Different measures of sample purity (immune infiltration, stromal component, or combined purity score) across breast cancer subsets. p53-only tumors (top right panel) show an increase in immune cell infiltration with p53 mutations from noCIN-like levels to acute CIN-like levels (consistent with our 3T3 data). MYC-only samples (grey boxplots in bottom row) show no change in infiltration depending on p53 status, and the CIN STAT1 ko tumors show a comparable infiltration level to noCIN tumors. **(c)** Linear associations of different predictors (x axis labels) with MYC gene expression or MYC Target expression (y panel) for either all samples (left panels) or stratified by p53 status (middle, right panels). MYC targets in p53 wild-type shows the strongest association with the STAT1 knockout signature, whereas there is no good predictor for p53 mutant tumors. **(d)** Type II ANOVA to quantify the relative importance of predictors. Each slice represents the level of MYC expression (top panels) and MYC Targets V1 (bottom panels) variance explained by one variable that cannot be explained by other variables. Taking into account multiple predictors, the STAT1 knockout signature explains more of the MYC target gene expression than MYC copy number, expression, or tumor subtype. The overall amount of variance (pie size) is highest in p53-mutant tumors (MYC; top panels) and in p53 wild-type tumors (MYC targets, bottom panels), respectively.

