## Supplementary figures and images for "Cancer tolerance to chromosomal instability is driven by Stat1 inactivation *in vivo*"

### Figure S1

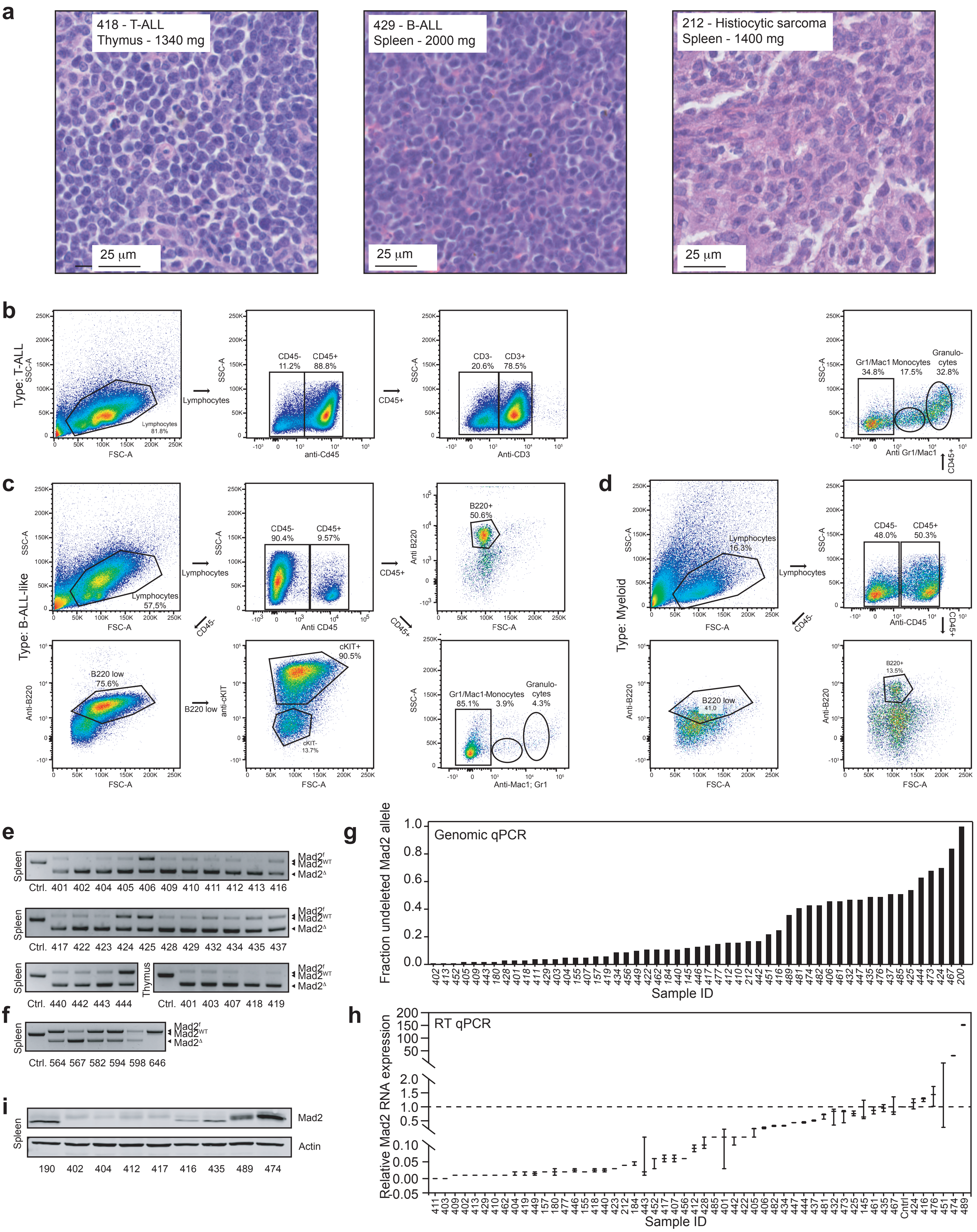


**Schubert *et al*, Figure S1**

### Figure S2

**Schubert *et al*, Figure S2**

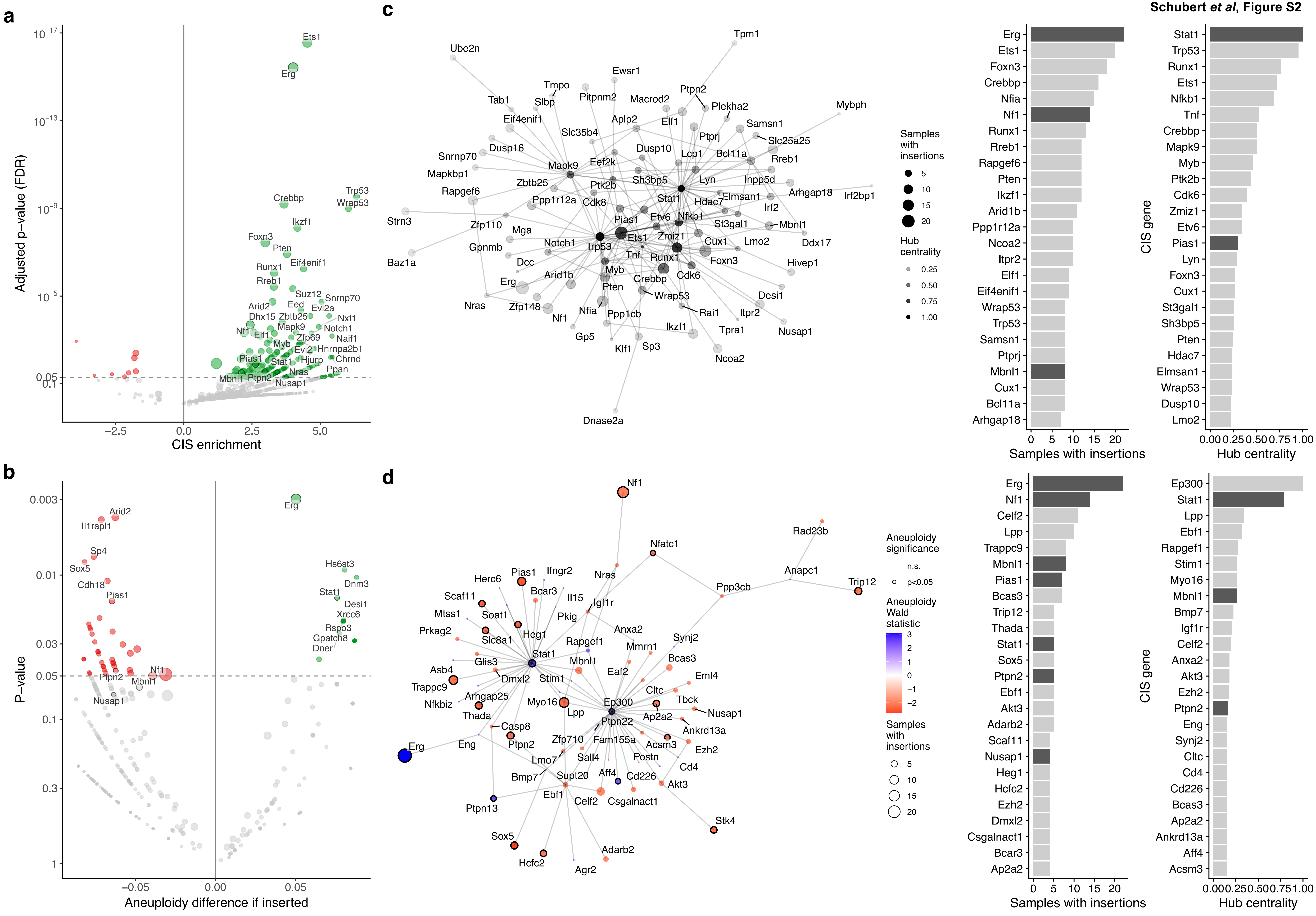

### Figure S3

**Schubert *et al*, Figure S3**

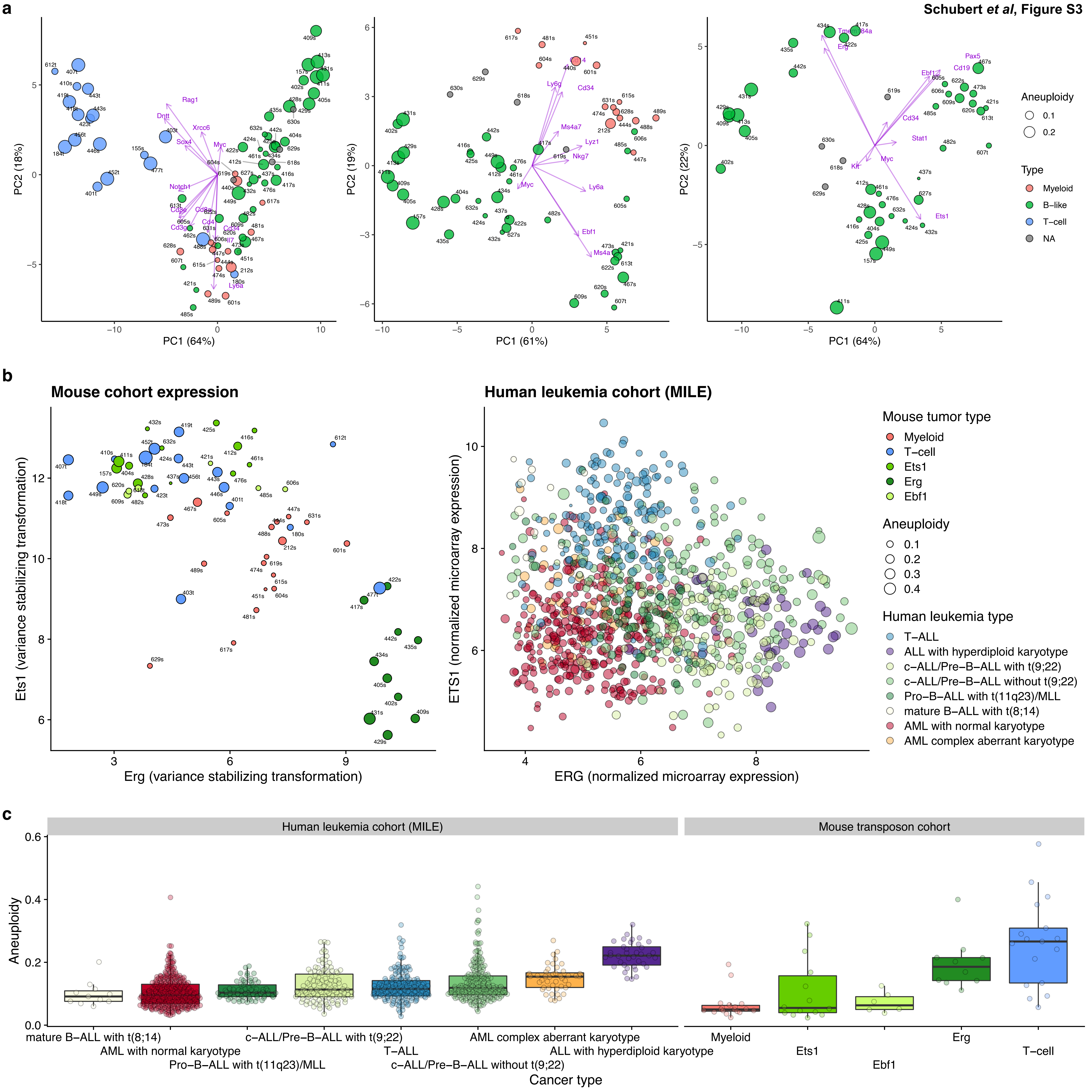

### Figure S4

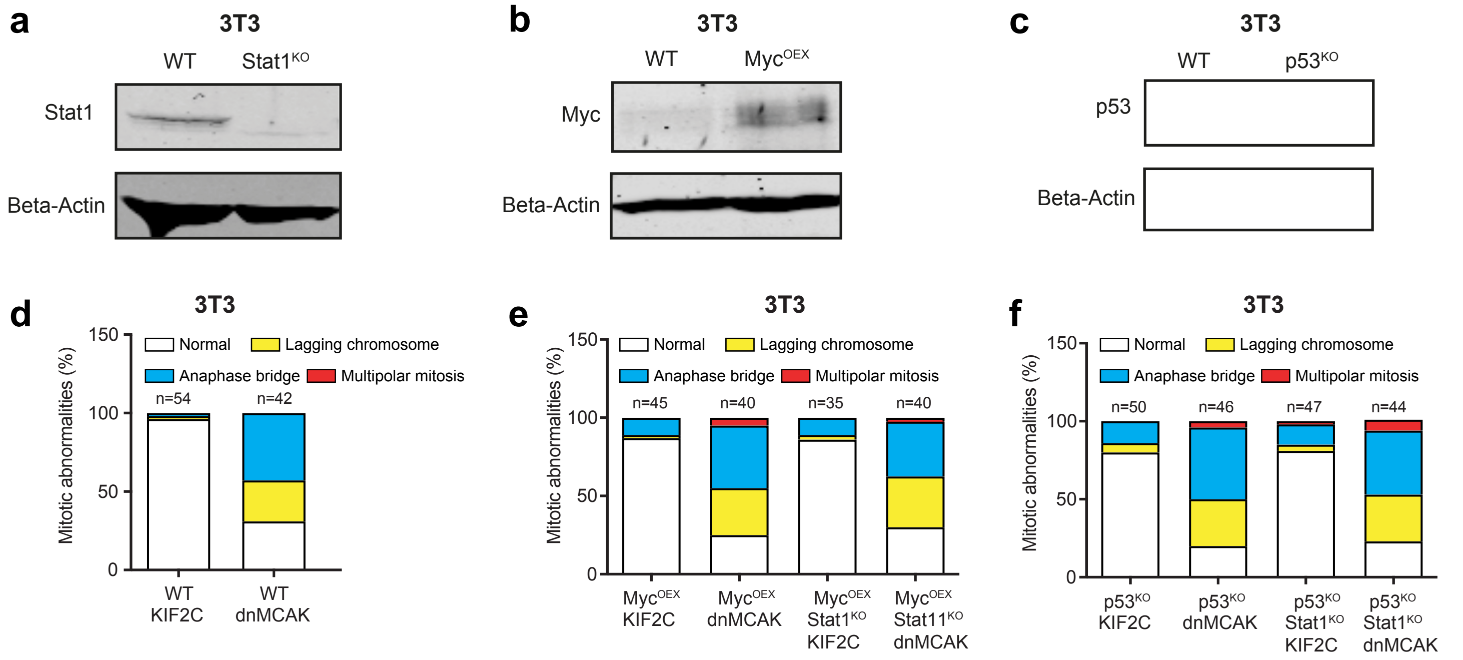


**Schubert *et al*, Figure S4**

### Figure S5

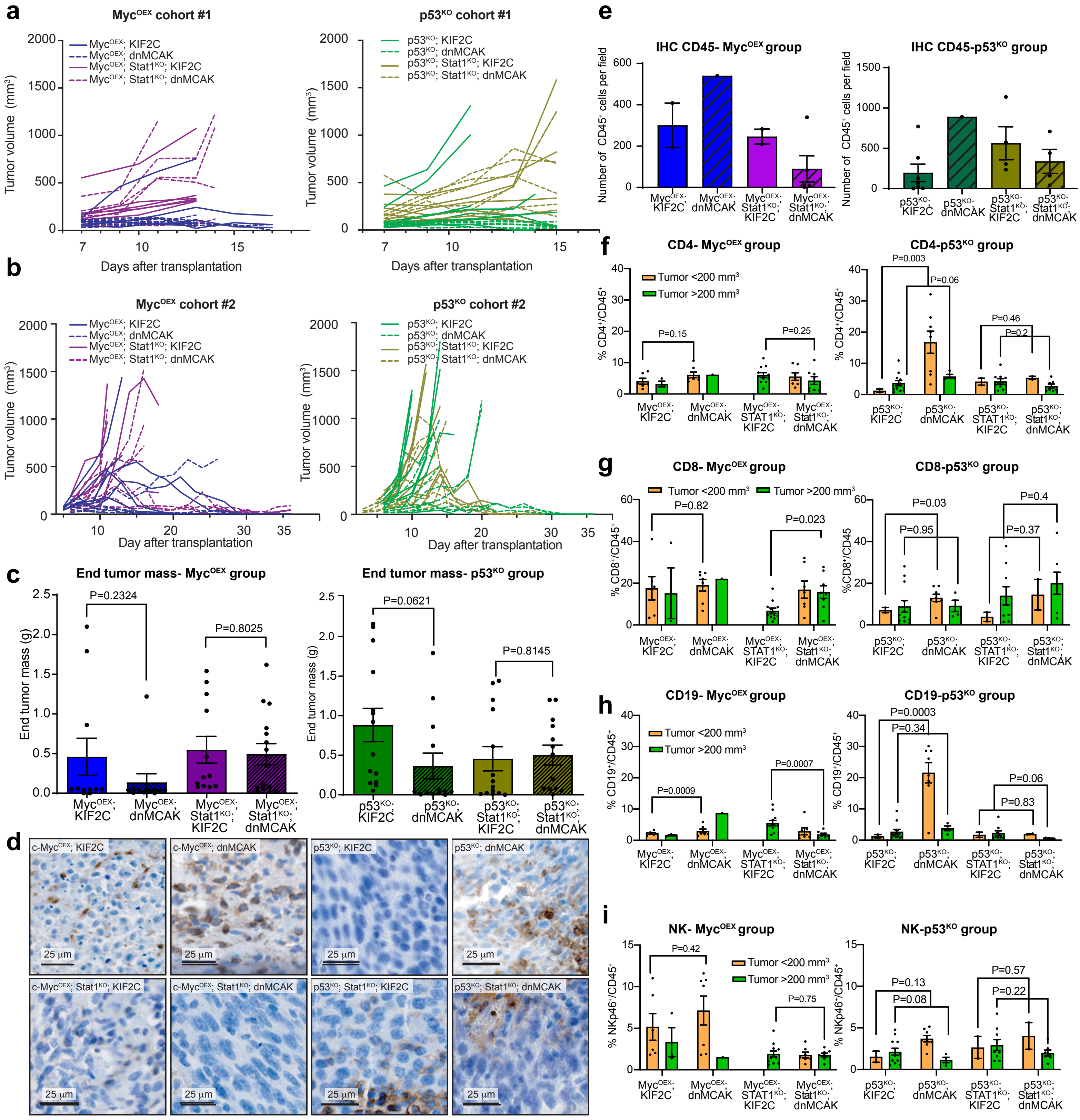


**Schubert *et al*, Figure S5**

### Figure S6

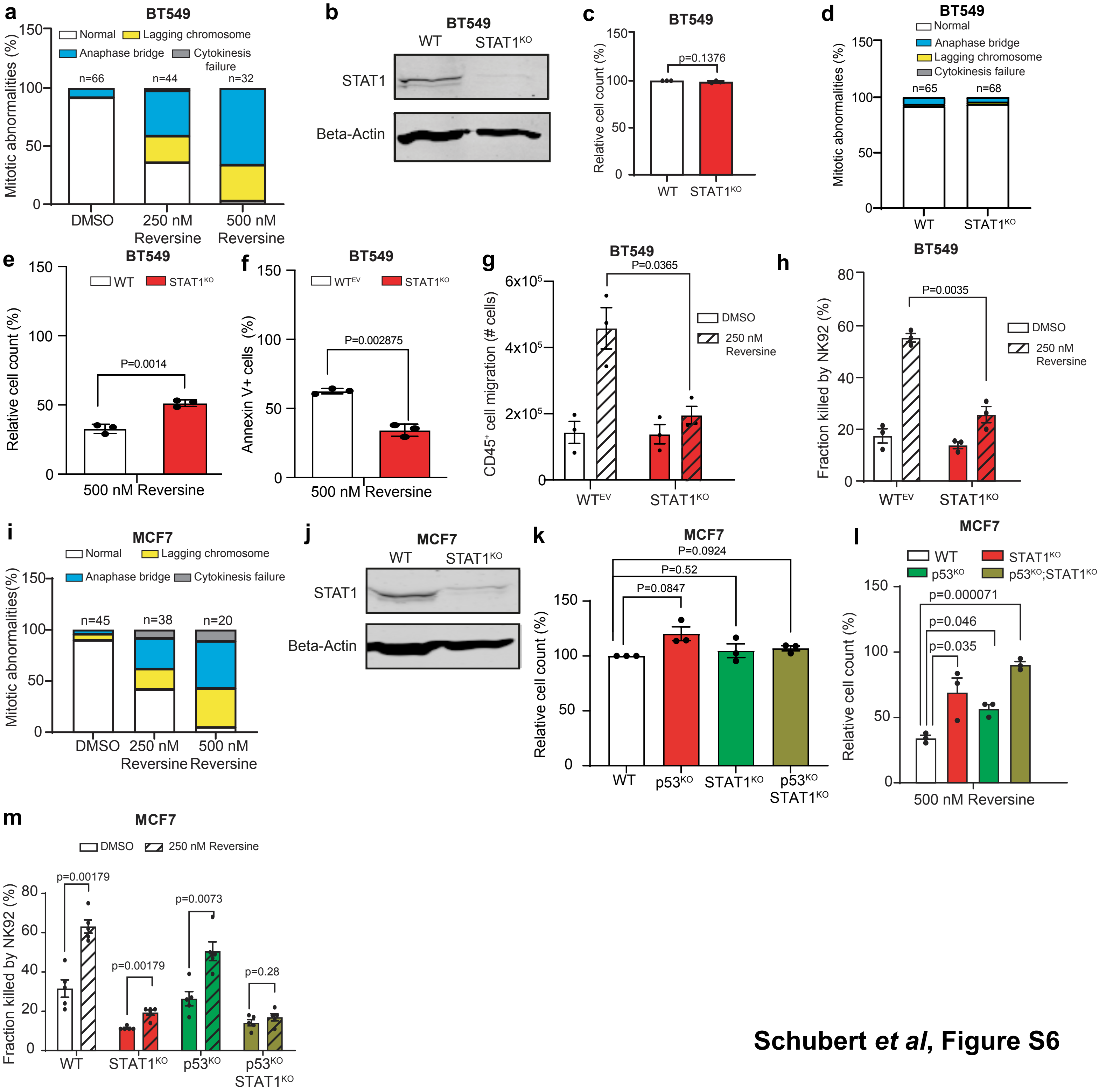


**Schubert *et al*, Figure S6**

### Figure S7

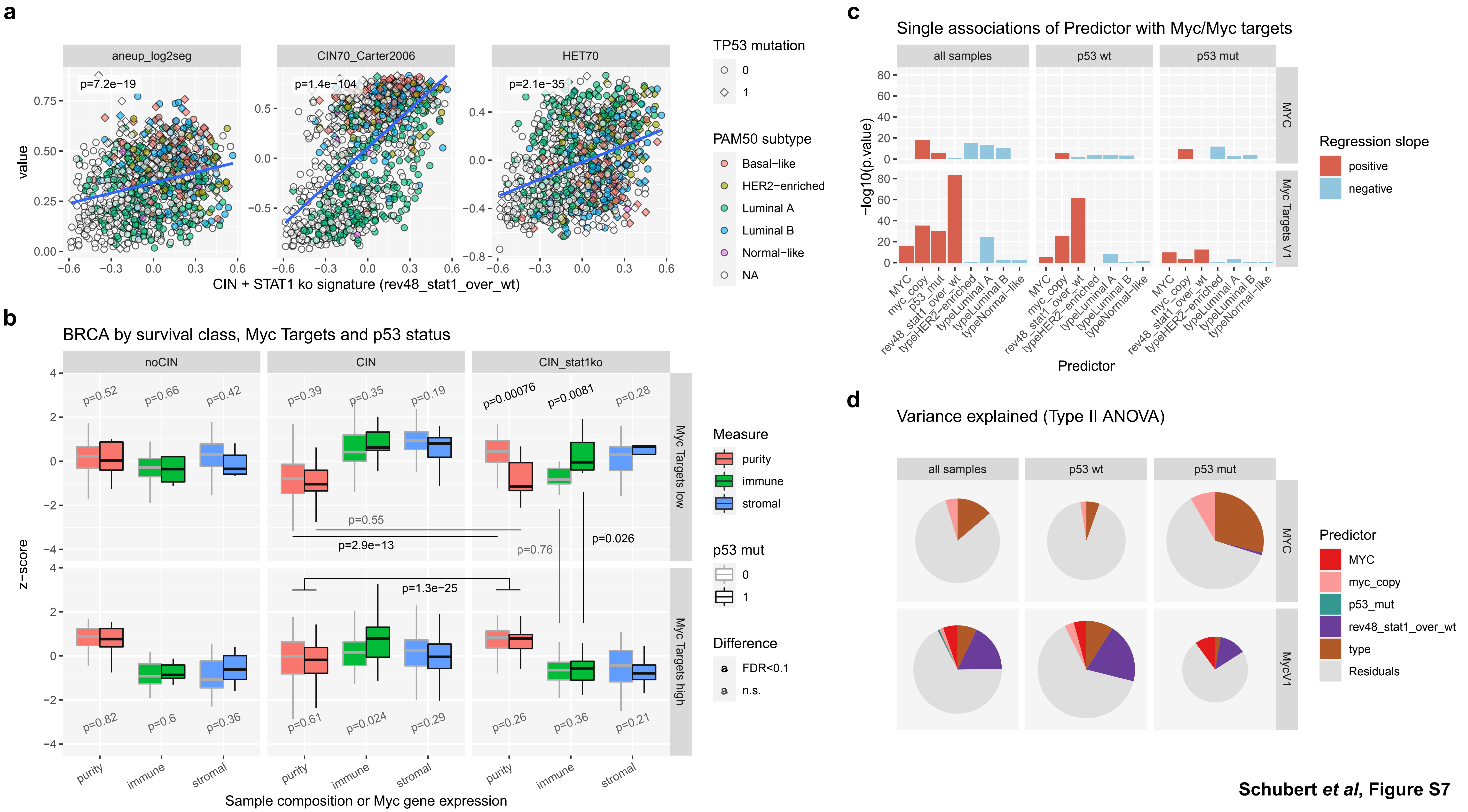


**Schubert *et al*, Figure S7**
